# predNMD: prediction of nonsense-mediated mRNA decay for improved clinical variant pathogenicity classification

**DOI:** 10.64898/2026.06.20.733449

**Authors:** Yaqi Su, Steven E Brenner

**Affiliations:** Department of Molecular and Cell Biology, University of California, Berkeley, CA 94720, USA; Center for Computational Biology, University of California, Berkeley, CA 94720, USA; Department of Plant and Microbial Biology, University of California, Berkeley, CA 94720, USA

## Abstract

The clinical consequence of a stop-gain variant depends on whether it ablates protein production by triggering nonsense-mediated mRNA decay (NMD) or yields truncated protein with residual, dominant-negative, or gain-of-function activity. This distinction is the key branch-point in applying PVS1, the strongest ACMG/AMP pathogenic evidence criterion. Current PVS1 guidelines and proposed SVC v4.0 successors risk unwarranted evidence assignment by employing only the 50nt rule for NMD prediction, which is central but incompletely captures NMD biology. We developed predNMD, a random forest classifier trained on 5,304 nonsense variants from GTEx, TCGA, GEUVADIS, and GREGoR. Feature selection reduced 166 candidates to 20 final features, 7 new to NMD prediction, including m6A density and TranslationAI. For variants predicted to not trigger NMD, predNMD infers the likely protein truncation. predNMD predictions aligned with deliberate clinical decisions where ClinGen Variant Curation Expert Panels (VCEPs) drew on evidence leading them to diverge from standard decision tree. predNMD agreed with VCEP on 6 of the 8 stop-gain variants for which they declined full PVS1, though the variants were predicted to trigger NMD by the 50nt rule and in loss-of-function disease genes. In leave-one-chromosome-out cross-validation predNMD reached AUC=0.79, and on independent test set outperformed 50nt rule (AUC 0.78 vs 0.65), nearly doubling discriminative signal above random (0.28 vs 0.15). predNMD likewise effectively discriminated NMD targets on BRCA1 and BARD1 saturation genome-editing data, where RNA abundance reflects NMD. These results support replacing the 50nt rule in clinical variant classification with predNMD, available as precomputed predictions covering all 13,968,776 possible stop-gain SNVs in GRCh38, installable codes, Docker image, and at predNMD.org.

## Introduction

Nonsense-mediated mRNA decay (NMD) is a conserved eukaryotic RNA surveillance pathway that detects and degrades transcripts containing premature termination codons (PTCs), thereby preventing the accumulation of truncated proteins with potentially deleterious effects. Genetic and molecular studies established that NMD is translation dependent and selectively targets mRNAs arising from nonsense variants, frameshifts, or errors in RNA processing^1–3^. Subsequent work demonstrated that NMD in eutherian mammals is coupled to pre-mRNA splicing and is influenced by exon-exon junction complexes (EJCs), which mark spliced transcripts and participate in distinguishing premature from normal translation termination events ^4–6^.

From a clinical perspective, NMD serves dual and sometimes opposing roles. On one hand, it acts as a quality-control mechanism that protects cells from dominant-negative, toxic, or gain-of-function effects of truncated proteins. Without NMD, many early stop-gain or frameshift variants would produce shortened proteins that retain partial domains and interfere with normal cellular functions ^7^. On the other hand, NMD can exacerbate disease by reducing functional gene dosage, leading to haploinsufficiency. ^8,9^ These contrasting effects make NMD a critical modifier of genotype-phenotype relationships in human disease and a key factor in the interpretation of predicted loss-of-function variants.

In mammals, the most widely used framework for predicting NMD activity is the exon junction complex–dependent “50nt rule,” which holds that a stop codon is recognized as premature if it lies more than approximately 50-55 nucleotides upstream of the final exon-exon junction. During the pioneer round of translation, ribosomes displace EJCs encountered within the coding sequence. If translation terminates upstream of a EJC, NMD is triggered, whereas termination within the last exon or sufficiently close to the final exon-exon junction generally permits transcript stability and normal protein production ^1,2,4–6^.

Despite its simplicity, the 50nt rule has proven highly informative and explanatory for clinical genetics. For example, Inoue et al., 2004 ^10^ showed that stop gained variants in SOX10 far enough upstream of normal stop codon tend to trigger NMD and lead to Waardenburg syndrome type 4 (WS4), a less severe phenotype consistent with haploinsufficiency. In contrast, nonsense variants that do not trigger NMD produce stable but deleterious truncated proteins, causing the far more severe PCWH syndrome (peripheral demyelinating neuropathy, central dysmyelinating leukodystrophy, Waardenburg syndrome, and Hirschsprung disease). Whether NMD is triggered thus determines not only disease severity but the type of syndrome that manifests.

A related principle underlies the long-recognized but mechanistically puzzling “frameshift rule” in dystrophinopathies. Many in-frame deletions of *DMD* result in Becker muscular dystrophy, a comparatively mild disorder, because partially functional dystrophin proteins missing internal domains are still produced. In contrast, smaller frameshift variants that sometimes are a subset of the Becker muscular dystrophy deletion causes Duchenne muscular dystrophy, a much more severe condition ^11–13^. This occurs because it often introduces upstream PTCs that trigger NMD, abolishing dystrophin expression. Experimental studies demonstrating phenotypic rescue in *mdx* mice through engineered C-terminal truncations further support the conclusion that NMD-mediated elimination of partially functional proteins can worsen disease outcomes ^14^.

Beyond its role in surveillance of pathogenic variants, NMD is also recognized as a widespread mechanism of normal gene regulation. Genome-wide analyses indicate that around one fifth of expressed human genes produce at least one transcript isoform that is constitutively degraded by NMD under physiological conditions. These transcripts are not aberrant byproducts but are generated through regulated RNA processing, most prominently alternative splicing. Alternative splicing coupled with NMD (called AS-NMD or RUST (regulated unproductive splicing and translation) ^15^) enables cells to modulate gene expression by controlling the balance between productive and unproductive isoforms, often through the inclusion or exclusion of a “poison exon” that introduces a PTC. Similarly, retention of introns within the 3’ UTR can deposit exon junction complexes downstream of the normal stop codon, causing it to be recognized as premature and thereby subjecting the transcript to NMD-mediated downregulation ^16^.

AS-NMD is particularly prevalent among genes encoding RNA-binding proteins and splicing factors, many of which auto-regulate or cross-regulate one another via feedback loops that route specific isoforms into the NMD pathway ^17^. This regulatory architecture stabilizes gene expression networks and buffers fluctuations in splicing factor abundance. Importantly, the same mechanisms can become pathogenic when dysregulated, especially in the context of noncoding or deep-intronic variants that alter splicing decisions.

An example is provided by *SCN1A*, where aberrant inclusion of a normally repressed poison exon leads to NMD-mediated loss of sodium channel expression and causes Dravet syndrome. Human genetic studies and mouse models have demonstrated that deep-intronic variants can activate this poison exon, triggering NMD and resulting in severe epileptic phenotypes ^18,19^. Notably, antisense oligonucleotide (ASO) therapies that promote poison exon exclusion can restore productive splicing and rescue gene expression, demonstrating that AS-NMD is a viable therapeutic target ^20^. Conversely, in certain dominant-negative or gain-of-function disorders, deliberately promoting poison exon inclusion or AS causing frameshift to induce NMD may be therapeutically beneficial, further underscoring the importance of accurately predicting NMD outcomes ^21^.

Comparative and evolutionary studies indicate that AS-NMD is an ancient and conserved mode of gene regulation. Many poison exons overlap ultraconserved elements, which are genomic regions exhibiting extraordinary sequence conservation across vertebrates, suggesting strong selective pressure to maintain NMD-based regulatory switches ^22^. Although the specific exon structures involved can vary, the coupling between alternative splicing and NMD is conserved from mammals to fungi, emphasizing its fundamental biological importance ^23^.

Given its profound impact on protein output, NMD plays a crucial role in clinical variant interpretation. In the ACMG/AMP guidelines, predicted loss-of-function variants such as nonsense and frameshift variants may receive very strong pathogenic evidence under the PVS1 criterion, the strongest code for pathogenic evidence. However, application of PVS1 explicitly depends on whether a variant is expected to trigger NMD ^24,25^. Variants that do not trigger NMD may produce truncated proteins with residual or altered function and therefore do not necessarily represent true loss-of-function alleles. Current PVS1 decision trees solely rely on a simplified assumption about NMD, i.e., inferred from the 50nt rule.

Accumulating evidence, however, demonstrates that NMD activity is influenced by numerous factors beyond EJC position, including proximity of the early stop codon to the start codon and the potential for translation reinitiation, length and composition of the 3′ UTR, number of downstream exons, local sequence context surrounding the early stop codon, RNA-binding protein occupancy, and stop-codon readthrough. Moreover, species-specific differences in NMD recognition mechanisms, such as a greater reliance on 3′ UTR length in flies and yeast, underscore the limitations of a single-rule framework for NMD prediction ^26–31^.

Several groups have attempted to model NMD activity using large-scale human genomic datasets. Rivas et al., 2015 ^32^ first introduced allele-specific expression as a proxy for NMD activity using GTEx data and trained a random forest classifier to distinguish NMD-triggering variants. Lindeboom et al.,2019 ^33^ developed NMDetective using TCGA data, identifying proximity to the start codon and unexpectedly long exons as important predictors for NMD activity. Teran et al., 2021 ^34^ applied LASSO regression to GTEx data and found that allele frequency, GC content, and evolutionary conservation contribute to NMD prediction, while Kim et al., 2024 ^35^ reported that gene-level intolerance to haploinsufficiency (LOEUF) predicts NMD outcomes. However, these models were generally trained on single data sources, achieved modest performance, were not released as broadly usable tools, and did not explicitly model downstream biological consequences when NMD does not occur, such as N-terminal truncation C-terminal truncation, or full-length protein production via translation readthrough.

Taken together, these limitations highlight a substantial gap between biological understanding of NMD and its practical use in clinical genomics. Reliance on the EJC-based 50nt rule alone risks misclassification of pathogenic variants, potentially delaying diagnosis or leading to inappropriate clinical decisions, and existing predictive models do not adequately capture uncertainty or biological diversity in NMD outcomes. Here, we address this gap by developing predNMD, a machine-learning method trained on integrated datasets from GTEx, TCGA, GEUVADIS, and GREGoR ^36–39^. Using systematic feature curation and selection, we identify a compact set of predictive features that improve NMD prediction accuracy. Our model achieves robust cross-validated performance, generalizes to independent datasets while outperforming previous NMD predictors available, and further provides biologically interpretable predictions, including the likely protein outcome when NMD does not occur. We explore how refined NMD prediction can improve application of the PVS1 criterion and ultimately enhance the classification of stop-gain variants for clinical diagnosis.

## Materials and Methods

### Training, validation, and test datasets

To train, validate, and test predNMD, we curated nonsense variants from seven independent sources that provide different proxies for NMD activity (Table 1). The training data included GREGoR (Genomics Research to Elucidate the Genetics of Rare Diseases) ^39^, a rare disease genomics cohort for which we quantified ASE from variant calls and RNA-seq data; GEUVADIS (Genetic European Variation in Disease) ^38^, a population-scale lymphoblastoid cell line resource with ASE measurements; GTEx (Genotype-Tissue Expression) ^36^, a large normal-tissue transcriptomic resource from which we derived ASE-based labels; and TCGA (The Cancer Genome Atlas) ^37^, a pan-cancer cohort for which we obtained previously processed transcript-level “NMD efficiency” estimates. Independent validation used the MMRF-TARGET dataset from Kim et al., 2024 ^35^, which combines processed “NMD efficiency” estimates derived from the Multiple Myeloma Research Foundation (MMRF) CoMMpass dataset and the Therapeutically Applicable Research to Generate Effective Treatments (TARGET) dataset. We further evaluated predNMD using BRCA1 and BARD1 saturation genome editing datasets with RNA abundance readouts for stop-gain mutations.

**Table 1.** Summary of datasets used for training, validating, and testing predNMD.

| Dataset | Use in predNMD | Data source / citation | NMD proxy | Binarization threshold | Number of nonsense variants |  |  |
| --- | --- | --- | --- | --- | --- | --- | --- |
|  |  |  |  |  | No-NMD | Indeterminate (excluded) | NMD-trigger |
| GREGoR | Training | RNA-seq BAM and VCF files from GREGoR Release 3 on AnVIL <sup>38</sup>, processed to ASE table following approach in Teran et al., 2021 <sup>33</sup> | Reference allele ratio | **NMD-trigger:** median reference allele ratio > 0.65 and at least half of high-ratio samples FDR-adjusted binomial p < 0.05.<br>**No-NMD:** median reference allele ratio < 0.55 and at least half of low-ratio samples FDR-adjusted p ≥ 0.05. | 615 | 901 | 421 |
| GEUVADIS | Training | GEUVADIS precomputed ASE table <sup>37</sup> | Reference allele ratio | Same as GREGoR | 219 | 308 | 248 |
| GTEX | Training | GTEX v8 ASE tables from AnVIL <sup>35</sup> | MODASE, calculated as 1 - posterior probability of no ASE <sup>33</sup> | **NMD-trigger:** median MODASE > 0.8 and at least half of high-MODASE samples FDR-adjusted binomial p < 0.05.<br>**No-NMD:** median MODASE < 0.2 and at least half of low-MODASE samples FDR-adjusted p ≥ 0.05. | 1,151 | 1,693 | 456 |
| TCGA | Training | Processed TCGA-derived <sup>36</sup> "NMD efficiency" data from Lindeboom et al., 2019 <sup>32</sup> | Transcript-level "NMD efficiency", defined as -log2 (expression of nonsense transcript / median expression of wild-type transcript) | **NMD-trigger:** "NMD efficiency" > 0.52.<br>**No-NMD:** "NMD efficiency" < 0.25. | 1,089 | 646 | 1,105 |
| MMRF-TARGET | Independent test set, evaluated only once | Processed “NMD efficiency” data from Kim et al., 2024 <sup>34</sup>, derived from MMRF <sup>44</sup> and TARGET <sup>45</sup> cohorts | ASE-derived “NMD efficiency”, defined as $-\log_2(\text{ASE\_RNA} / \text{ASE\_DNA})$ | **NMD-trigger:** “NMD efficiency” $> -\log_2((1 - 0.65) / 0.5) \approx 0.515$.<br>**No-NMD:** “NMD efficiency” $< -\log_2((1 - 0.55) / 0.5) \approx 0.152$. | 158 | 219 | 161 |
| BRCA1 MAVE | Experimental evaluation | BRCA1 saturation genome editing data from Findlay et al., 2018 <sup>42</sup> | RNA score from cDNA/genomic DNA variant frequency normalization | **NMD-trigger:** RNA score $< -1$ (~50% reduction in mRNA)<br>**No-NMD:** RNA score $\geq -1$ | 41 | NA | 89 |
| BARD1 MAVE | Experimental evaluation | BARD1 saturation genome editing data from Woo et al., 2025 <sup>43</sup> | RNA score from cDNA/genomic DNA variant frequency normalization | Same as BRCA1 MAVE | 98 | NA | 220 |
| BRCA1 mutations in Perrin-Vidoz’s assay overlapping Findlay’s assay | Experimental evaluation | NMD status reported in Perrin-Vidoz et al., 2002 <sup>46</sup> for nonsense mutations also in Findlay’s BRCA1 MAVE dataset <sup>42</sup> | Relative steady-state abundance of the transcript carrying nonsense variant versus the wild-type in patient-derived lymphoblastoid cell lines | Perrin-Vidoz et al categorical calls <sup>46</sup> | 6 | NA | 1 |

For the GREGoR dataset, we obtained 558 matched VCF files and aligned RNA-seq BAM files from Release 3 on the AnVIL platform. Each VCF was annotated using Ensembl Variant Effect Predictor (VEP) v104 ^40^ to identify heterozygous nonsense variants, and allele-specific expression (ASE) was measured using GATK ASEReadCounter ^41^ with minimum read depth of 10, mapping quality of 20, and base quality of 20. We performed binomial tests at each variant site and applied FDR correction to assess allelic imbalance. Variants were classified as “NMD-trigger” if they exhibited median reference allele ratio > 0.65 across all samples carrying that variant, and among the subset of samples with ratio > 0.65, at least half also showed FDR-corrected p-value < 0.05. Variants were classified as “no-NMD” if they showed median reference allele ratio < 0.55, and among the subset of samples with ratio < 0.55, at least half also showed FDR-corrected p-value ≥ 0.05. Variants not meeting either criterion were excluded, yielding 1,036 unique variants. For the GEUVADIS dataset, we downloaded merged variant files with pre-computed ASE records from the European Bioinformatics Institute (https://www.ebi.ac.uk/biostudies/files/E-GEUV-1/E-GEUV-1/analysis_results/GD462.ASE.COV8.ANNOT_PTV.txt.gz, accessed on 12/12/2024) and applied the same classification criteria, obtaining 467 unique variants.

The GTEx v8 dataset required a modified approach to account for tissue-level variation in “NMD efficiency”. We downloaded all pre-processed ASE tables from AnVIL and calculated the probability of ASE (MODASE) for each variant following Teran et al., 2021 ^34^. MODASE is derived from a Bayesian statistical framework ^32^ that leverages the multi-tissue design of GTEx to predict the probability of allele-specific expression at a variant site. Specifically, MODASE is calculated as 1 minus the posterior probability of no ASE, integrating information from multiple observations of a variant across tissues within the same individual to reduce noise from tissue-specific variations and technical artifacts. After calculating MODASE values and performing binomial tests (based on ASE) at each variant site, we classified variants using the criteria as follows: variants were classified as “NMD-trigger” if they exhibited median MODASE > 0.8 across all samples carrying that variant, and among the subset of samples with MODASE > 0.8, at least half also showed FDR-corrected p-value < 0.05. Variants were classified as “no-NMD” if they showed median MODASE < 0.2, and among the subset of samples with MODASE < 0.2, at least half also showed FDR-corrected p-value ≥ 0.05. Variants not meeting either criterion were excluded, yielding 1,607 unique variants.

For the TCGA dataset, we obtained pre-processed data directly from Dr. Rik Lindeboom corresponding to their published study ^33^. Their dataset quantifies “NMD efficiency” as −log₂(expression of nonsense transcript / median expression of wild-type transcript), measuring the degree of transcript degradation for each nonsense variant. We applied their published thresholds: “NMD efficiency” > 0.52 for “NMD-trigger” variants, < 0.25 for “no-NMD” variants, excluding intermediate values, which identified 2,194 unique variants. Therefore, there were a total of 5304 high-quality variants in the training dataset which was used to train the Random Forest model for NMD prediction.

For the independent MMRF-TARGET validation dataset, we directly obtained processed data from Supplementary Data 12 from Kim et al., 2024 ^35^, which defines “NMD efficiency” as −log₂(ASE_RNA/ASE_DNA) to normalize RNA-level allelic imbalance against DNA-level imbalance, thereby controlling for germline copy number variations and somatic alterations. To maintain consistency with our ASE-based classification criteria from GREGoR and GTEx, we applied analogous thresholds: variants with “NMD efficiency” > −log₂((1−0.65)/0.5) ≈ 0.515 were classified as “NMD-trigger”, those with “NMD efficiency” < −log₂((1−0.55)/0.5) ≈ 0.152 were classified as “no NMD”, and intermediate variants were excluded, yielding 319 variants.

The BRCA1 MAVE (multiplexed assay of variant effect) dataset was obtained from MAVEdb ^42^ (score set urn:mavedb:00000097-0-2), originally published by Findlay et al., 2018 ^43^. The BARD1 MAVE dataset was obtained from a saturation genome editing study by Woo et al., 2025 ^44^. Both datasets employed saturation genome editing (SGE) to systematically assess single nucleotide variants for effects on cell survival and RNA abundance. To assess the impact of variants on mRNA expression and potential degradation by NMD, we utilized the RNA scores from each study, which quantify mRNA expression levels by normalizing variant frequencies in cDNA to their frequencies in genomic DNA. This normalization accounts for differences in editing efficiency, isolating the effect of each variant on mRNA abundance and allowing RNA scores to reflect post-transcriptional regulatory effects including NMD. We restricted downstream analyses to stop-gained single-nucleotide variants with available RNA scores.

For variants overlapping multiple transcript isoforms in the GEUVADIS, GTEx, and GREGoR datasets, we assigned each variant to a single representative transcript using hierarchical prioritization. We first prioritized the canonical transcript annotated by VEP, which identifies the most functionally relevant transcript for each gene. When no canonical transcript was available or when multiple canonical transcripts existed, we selected the longest transcript isoform to maximize the likelihood of capturing complete coding sequence features. The TCGA and MMRF-TARGET datasets were already de-duplicated with one unique transcript per variant. All distance-based and transcript structure features were calculated using genome annotations from Ensembl GRCh37 (release 87) for GEUVADIS and TCGA, Ensembl GRCh38 (release 104) for GTEx, MMRF-TARGET, and MAVE datasets, and GENCODE v41 for GREGoR.

In this study, we define “full PVS1” as the exact evidence code “PVS1”. To assess concordance between predNMD predictions and expert clinical variant interpretation, we curated stop-gain variants from the ClinGen Evidence Repository (https://erepo.clinicalgenome.org/evrepo/, accessed on 7/25/2025) that were not assigned full PVS1. We examined the evidence codes assigned by Variant Curation Expert Panels and considered variants assigned reduced-strength PVS1 codes, including “PVS1_Strong”, “PVS1_Moderate”, or “PVS1_Supporting”, as not assigned full PVS1. This distinction reflects the PVS1 decision framework, in which stop-gain variants predicted to trigger NMD may receive full PVS1, whereas variants predicted not to trigger NMD may receive reduced PVS1 strength at most. We then compared predNMD predictions for this curated set with the expert-panel rationale for not applying full PVS1. The RUNX1 c.356dup (p.Ala120GlyfsTer18) variant was originally included but later removed after the Myeloid Malignancy VCEP confirmed its curation had incorrectly assumed the variant does not undergo NMD; the panel agreed the early stop codon is upstream of the last exon–exon junction and predicted to undergo NMD

### Random forest classification

We implemented predNMD as a random forest classifier using the RandomForestClassifier class from scikit-learn v1.6.0 ^45^ in Python 3.10. Hyperparameter optimization was performed using randomized search (RandomizedSearchCV) with 100 iterations to efficiently explore the parameter space, employing a leave-one-chromosome-out cross-validation (LOCO-CV) strategy. In LOCO-CV, variants from each chromosome are iteratively held out as a validation set while training on the remaining chromosomes, ensuring that model performance estimates reflect the ability to generalize to variants in genomic regions not seen during training and accounting for potential chromosome-specific biases in variant properties or annotation quality. The search was optimized for the area under the receiver operating characteristic curve (AUC-ROC).

The hyperparameter search space included n_estimators (number of decision trees: 50, 100, 200, 300, 500, 800, 1000), max_depth (maximum tree depth: 3, 5, 10, 15, 20, 25, 30, None), min_samples_split (minimum samples required to split an internal node: 2, 5, 10, 15, 20), min_samples_leaf (minimum samples required at leaf nodes: 1, 2, 4, 8, 12), max_features (number of features considered at each split: ‘sqrt’, ‘log2’, None, 0.3, 0.5, 0.7), bootstrap (whether to use bootstrap sampling: True, False), class_weight (weights to address class imbalance: None, ‘balanced’, ‘balanced_subsample’), and max_samples (fraction of samples to draw for each tree when bootstrap=True: None, 0.7, 0.8, 0.9). The optimal hyperparameters selected were: n_estimators=1000, max_depth=25, min_samples_split=2, min_samples_leaf=8, max_features=’sqrt’, class_weight=’balanced’, bootstrap=True, and max_samples=0.7.

We quantified feature importance using the built-in random forest feature importance based on mean decrease in impurity (MDI), which measures the average reduction in Gini impurity when a feature is used to split nodes across all trees in the forest. This metric provides a measure of each feature’s contribution to the model’s predictive performance, with higher values indicating features that are more important for distinguishing between NMD-trigger and no-NMD variants.

### Feature Selection

We first curated 166 computable candidate features that could plausibly contribute to NMD prediction. These features were selected to capture biological factors previously implicated in NMD activity, transcript and variant contexts that define the relationship between an early stop codon and exon-exon junctions, and translation-related contexts that may influence whether a transcript undergoes degradation or produces a truncated protein. The candidate features were organized into 10 broad categories: position- and distance-related features, transcript structure features, sequence element features, m6A-related features, translation readthrough-related features, translation reinitiation-related features, gene characteristics, allele frequency, conservation at the variant site, and translation efficiency-related features. Detailed definitions, data sources, and calculation procedures for all 166 candidate features are provided in the Supplementary Description.

We then performed feature selection using the Boruta algorithm ^46^, an iterative wrapper method that identifies all relevant features by creating shadow features, defined as randomized copies of the original features, and comparing the importance of real features against these random probes. Features that consistently demonstrate higher importance than the best shadow features are retained as confirmed features, while features performing no better than random are rejected. We implemented Boruta using the BorutaPy package with max_iter=200, alpha=0.05 (significance level), two_step=True (enabling a two-stage selection process to better handle borderline features), and n_estimators=’auto’. The base estimator within Boruta was a RandomForestClassifier with 300 trees, max_depth=20, min_samples_split=5, and min_samples_leaf=2 to ensure stable feature importance rankings during the selection process. Boruta identified 20 confirmed features with no tentative features, indicating strong and unambiguous evidence for the relevance of these features to NMD prediction.

The 20 selected features are summarized in Table 2. These features were curated from multiple genomic databases and calculated using standardized bioinformatics approaches. Gene expression levels were quantified as mean transcripts per million (TPM) across all GTEx v10 tissues with TPM > 0. N⁶-methyladenosine (m6A) modification sites were obtained from m6A-Atlas v2.0 ^47^, using only high-resolution data to ensure mapping accuracy. The Codon Adaptation Index (CAI) was calculated for the 25-codon window upstream of the early stop codon as the geometric mean of relative codon usage frequencies, using the human codon usage table from the HIVE-CUTs database ^48^. Specifically, CAI was computed as exp[(1/L) × Σ_i_ ln(w_i_)], where L is the number of codons and w_i_ is the relative adaptedness value for codon i defined as the frequency of codon i divided by the frequency of the most common codon for the same amino acid ^49^. We calculated the CAI difference by subtracting the CAI of the 25-codon window upstream of the early stop codon from the CAI of the corresponding region relative to the normal stop codon, capturing local changes in codon optimality that may affect translation efficiency. Gene-level intolerance to haploinsufficiency was measured using the loss-of-function observed/expected upper bound fraction (LOEUF) from gnomAD v4.1 constraint metrics ^50^. Evolutionary conservation was assessed using phyloP scores extracted from 100-way vertebrate alignments available through the UCSC Genome Browser ^51^. Translation initiation site scores given by TranslationAI (v1.0) at the first in-frame AUG codon downstream to the early stop codon were used to assess reinitiation potential, while translation termination site scores given by TranslationAI at the early stop codon were used to quantify termination signal strength ^52^. Population allele frequencies were extracted from gnomAD, using gnomAD v4.1 for variants called with GRCh38/hg38 assembly (GTEx, GREGoR, MMRF-TARGET, and MAVE datasets), and gnomAD v2.1 for variants called with GRCh37/hg19 assembly (GEUVADIS and TCGA). All the other transcript structure related features were calculated using reference genome annotations from Ensembl (Ensembl GRCh37 release 87 for GEUVADIS and TCGA; Ensembl GRCh38 release 104 for GTEx, MMRF-TARGET, and MAVE) and GENCODE (GENCODE v41 for GREGoR).

**Table 2:** Features used in predNMD.

| Feature name | Type | Description | Feature category | Experimentally validated | Mechanistically sensible | New in predNMD |
| --- | --- | --- | --- | --- | --- | --- |
| ETC < 50 nt upstream of last exon-exon junction | Categorical | Whether the early stop codon is located < 50 nt (True) or ≥ 50 nt (False) upstream of the last exon-exon junction | Transcript structure | Yes <sup>2</sup> | Yes | No |
| Number of downstream exons | Integer | Number of exons downstream of the early stop codon | Transcript structure | Yes <sup>29</sup> | Yes | No |
| Distance to ETC-containing exon end | Integer | Distance of the early stop codon to the end of the exon containing the early stop codon | Transcript structure | No | Yes | No |
| Length of ETC-containing exon | Integer | Length of the exon containing the early stop codon | Transcript structure | No | Yes | No |
| Distance to the NTC | Integer | Distance of the early stop codon to the normal stop codon of the transcript | Transcript structure | No | Yes | No |
| CDS position of the ETC | Integer | Position of the early stop codon at the CDS region | Translation related | Yes <sup>60</sup> | Yes | No |
| TranslationAI score of ETC | Continuous | TSS (Translation Termination Site) score given by TranslationAI at the early stop codon site | Translation related | No | Yes | Yes |
| Mean gene expression | Continuous | Mean expression level (TPM) of the gene containing the early stop codon curated from GTEx | Gene level | No | Yes | No |
| TranslationAI score of downstream in-frame AUG | Continuous | TIS (Translation Initiation Site) score given by TranslationAI at the first in-frame AUG downstream of the early stop codon | Translation related | No | Yes | Yes |
| m6A density upstream of ETC (CDS region) | Continuous | m6A density (m6A count/distance between CDS start and the early stop codon) of the CDS region | RNA modification | No | Yes | Yes |
| GC content | Continuous | GC content of the transcript containing the early stop codon | Sequence context | Yes <sup>61</sup> | Yes | No |
| Distance to 3' UTR end | Integer | Distance of the early stop codon to the end of 3' UTR | Transcript structure | Yes <sup>26</sup> | Yes | No |
| phyloP | Continuous | Evolutionary conservation score at the early stop codon site from the 100-way vertebrate multiple sequence alignment | Evolution related | No | No | No |
| m6A density of whole transcript | Continuous | m6A density (m6A count/transcript length) of the whole transcript | RNA modification | No | Yes | Yes |
| CAI difference 25 codons upstream of ETC & NTC | Continuous | Codon Adaptation Index difference between 25 codons upstream of the early stop codon and that of the normal stop codon | Sequence context | No | Yes | Yes |
| LOEUF | Continuous | Loss-of-function observed/expected upper bound fraction from gnomAD | Gene-level | No | Maybe | No |
| AF | Continuous | Allele frequency of the variant from gnomAD | Population genetics | No | No | No |
| Distance to first in-frame AUG | Integer | Distance of the early stop codon to the first in-frame AUG downstream of the early stop codon | Translation related | No | Yes | Yes |
| Distance to first out-of-frame AUG | Integer | Distance of the early stop codon to the first out-of-frame AUG downstream of the early stop codon | Translation related | No | Yes | Yes |
| Number of upstream exons | Integer | Number of exons upstream of the early stop codon | Transcript structure | Yes <sup>29</sup> | Yes | No |

\* **ETC**: Early termination codon. **NTC**: Normal termination codon.
\* **Mechanistically sensible**: Whether the biological property underlying the feature is something a cell could plausibly sense and respond to through a molecular mechanism, as opposed to a population- or evolution-level summary external to any cell.

### SHAP analysis for predicting protein truncation outcome

When NMD is not triggered, an early termination codon may lead to production of either an N-terminally truncated protein through downstream reinitiation or a C-terminally truncated protein through premature termination. To estimate which truncated protein outcome was more likely for each variant predicted as not triggering NMD, we applied SHapley Additive exPlanations (SHAP) analysis ^53^ to quantify the contribution of different feature groups to the model prediction.

We used the TreeExplainer implementation from the SHAP Python library ^54^, which provides efficient exact calculations for tree-based models by leveraging their internal structure. Features were assigned to three biologically motivated groups: N-terminal features, representing evidence related to downstream translation reinitiation; C-terminal features, representing evidence related to transcript contexts compatible with not triggering NMD and production of a C-terminally truncated protein; and general features, representing sequence, evolutionary, expression, and transcript-level properties that may affect NMD prediction but do not uniquely distinguish between N- and C-terminal protein outcomes.

To estimate the relative probability of N- versus C-terminal truncation outcomes, we first summarized the SHAP values within each feature group. For each variant, we calculated category-level SHAP contribution scores by summing the SHAP values of all features within each category. We denote these summed scores as S_N, S_C, and S_G, representing the total SHAP contribution from the N-terminal, C-terminal, and general feature groups, respectively.

Because SHAP values are additive contributions to the model output, these summed scores represent the overall contribution of each feature group to the predicted NMD-trigger probability. For variants predicted not to trigger NMD, defined as predNMD NMD-trigger probability < 0.5, we converted the N-terminal and C-terminal category-level SHAP contribution scores into relative protein-outcome probabilities using a two-class softmax ^55^. Softmax was used because it transforms the two outcome scores into normalized probabilities that sum to one while preserving their relative support. Because negative SHAP values decrease the predicted NMD-trigger probability, a more negative category-level SHAP score was interpreted as stronger support for the corresponding truncated protein outcome. The relative protein-outcome probabilities were calculated as:

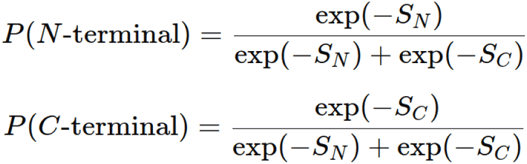

The general contribution score S_G was not included in the final softmax because it contributes equally to both protein-outcome scores and therefore cancels during normalization.

This transformation assigns higher probability to the protein outcome whose feature category contributes more strongly toward reduced NMD-trigger probability. If one of S_N or S_C was negative while the other was positive, the outcome corresponding to the negative score was selected as the supported truncation outcome, because only that feature category contributed toward reduced NMD-trigger probability. In cases where both S_N and S_C were positive, indicating that both N-terminal and C-terminal feature categories increased rather than decreased the predicted NMD-trigger probability, we assigned equal probabilities of 0.5 to reflect mechanistic uncertainty. This approach provides a relative estimate of whether an N-terminally or C-terminally truncated protein is more likely to be produced when a variant is predicted not to trigger NMD.

### Implementations of predNMD

The predNMD method is implemented as an open-source Python software package and is distributed through multiple access modalities to facilitate broad use in research and clinical genomics workflows. The core predNMD package can be installed from its public GitHub repository (https://github.com/BrennerLab/predNMD). To ensure reproducibility and simplify dependency management, we also provide a pre-built Docker image (https://hub.docker.com/r/brennerlab/prednmd) that encapsulates the complete runtime environment and reference resources required for prediction. This containerized implementation enables execution of predNMD in heterogeneous computing environments without local software configuration.

In addition to local execution, we provide precomputed predNMD predictions for all possible stop-gain single-nucleotide variants: 13,968,776 on Ensembl GRCh38 (release 104) and 12,714,784 on GRCh37 (release 87). These predictions can be accessed through the predNMD web interface at predNMD.org for single-variant lookup or queried in batch using precomputed tables and a lightweight command-line script. This allows users to rapidly annotate either individual variants or large VCF files without running the full predNMD prediction pipeline.

Together, the installable package, Docker image, web interface, and precomputed predictions allow users to access predNMD either through local execution, single-variant lookup, or batch processing of large variant files, enabling straightforward integration of predNMD predictions into clinical variant interpretation workflows.

## Results

### Feature selection identified 20 predictive features for predNMD

We initially compiled 166 candidate features involving transcript structure, sequence context, evolutionary conservation, translation signals, RNA modifications, and population genetics metrics for predicting NMD outcomes (Supplementary Figure 1). Preliminary analysis revealed that many of these features exhibited extremely low random forest importance scores, suggesting they contribute noise or redundancy rather than true predictive signal. To identify a parsimonious set of informative features and reduce the risk of overfitting, we applied the Boruta algorithm, an iterative wrapper method that compares feature importance against randomized shadow features to statistically distinguish relevant predictors from noise ^46^. Boruta identified 20 confirmed features and no tentative features, indicating strong and unambiguous evidence for the relevance of these features to NMD prediction.

The selected features span multiple biological categories (Table 2). Transcript structure features include the canonical 50nt rule, number of downstream and upstream exons, distance from the early stop codon to the exon end, length of the exon containing the early stop codon, distance to the normal termination codon, and distance to the end of 3’ UTR. Translation-related features include CDS position of the early stop codon, distances from the early stop codon to the first downstream in-frame and out-of-frame AUG codons, and two scores derived from TranslationAI ^52^, a deep learning model trained to predict translation initiation and termination efficiency from sequence context. In particular, we used TranslationAI’s translation initiation site (TIS) score at the first in-frame AUG codon downstream of the early stop codon to quantify reinitiation potential, and the translation termination site (TSS) score at the early stop codon to assess termination signal strength. RNA modification features include m6A density across the transcript and also specifically in the CDS region upstream of the early stop codon. Sequence features include GC content and CAI difference between the 25-codon window upstream of the early stop codon versus the normal stop codon. Gene-level features include mean expression level and LOEUF score, while population genetics is represented by gnomAD allele frequency and evolutionary feature is reflected by phyloP conservation score. Of these 20 features, seven represent novel predictors not previously incorporated into published NMD prediction models, while the remaining 13 have been used in previous models, either with established biological mechanisms or as empirically identified predictors without clear mechanistic interpretation (Figure 1A).

**Figure 1.**
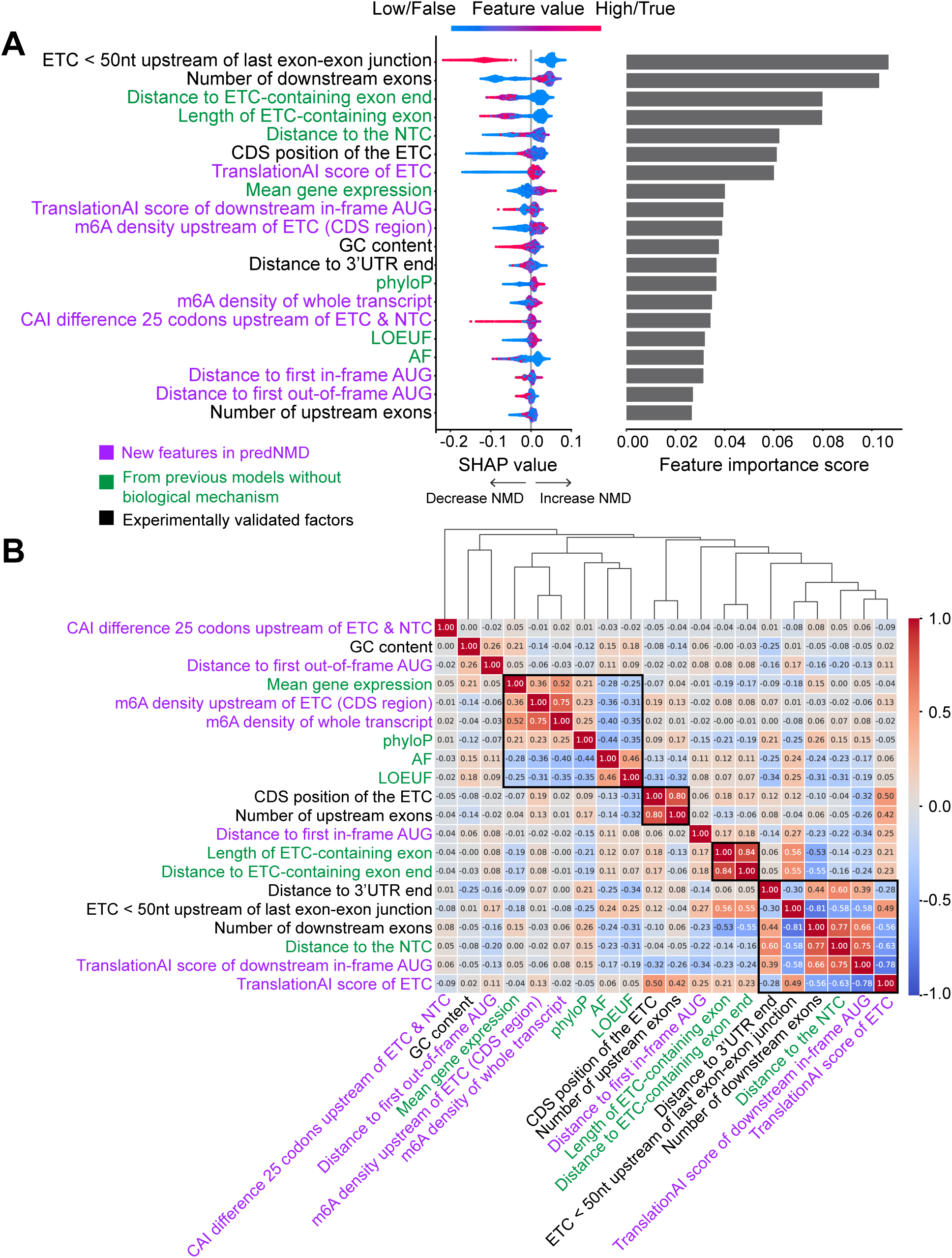
Importance and correlation analysis of features included in predNMD. (A) Feature importance of predNMD features. Left: SHAP beeswarm plot showing feature contributions to individual predictions, where x-axis position indicates impact on NMD probability (positive SHAP value is associated with increasing NMD-trigger probability and vice versa). The color of the point indicates feature values (red = high/True and blue = low/False for continuous/categorical features). Right: Mean Decrease Impurity (MDI) scores from the Random Forest model. (B) Spearman correlation matrix with hierarchical clustering based on absolute correlation value. Color intensity indicates correlation strength (red = positive, blue = negative).

Using mean decrease in impurity (MDI) from the random forest model, we quantified the relative importance of each feature in predNMD predictions (Figure 1A). The 50nt rule feature ranked first by MDI. In the SHAP analysis, high/true values for this feature, corresponding to early stop codons located within 50 nt upstream of the last exon-exon junction, were associated with negative SHAP values and therefore lower predicted NMD-trigger probability. Conversely, low/false values, corresponding to early stop codons located farther than 50 nt upstream of the last exon-exon junction, were associated with positive SHAP values and higher predicted NMD-trigger probability. This pattern is consistent with the canonical 50nt positional rule for mammalian NMD ^2^. The number of downstream exons ranked second by MDI. In the SHAP analysis, higher numbers of downstream exons were associated with positive SHAP values, indicating higher predicted NMD-trigger probability, whereas lower numbers of downstream exons were associated with negative SHAP values. This association is consistent with EJC-dependent models of NMD, in which exon-exon junctions downstream of an early stop codon can provide potential sites for EJC-associated NMD signaling ^56,57^. Distance to the end of the exon containing early stop codon and length of the exon containing early stop codon were also among the most important features. These features were previously identified as significant predictors by Lindeboom et al., 2019 ^33^ and Teran et al., 2021 ^34^, but without a clear biological mechanism to explain their contribution. We offer a potential mechanistic explanation for these observations, which will be explored in the Discussion.

Among the novel features in predNMD, the two TranslationAI-derived scores showed substantial predictive importance and may act in concert to determine NMD outcomes (Figure 1A). The TranslationAI score at the early stop codon captures the effectiveness of translation termination, while the TranslationAI score at the downstream in-frame AUG quantifies reinitiation potential. SHAP analysis showed that lower TranslationAI termination scores at the early stop codon were associated with reduced predicted NMD-trigger probability. This association is consistent with models in which inefficient termination or stop-codon readthrough can reduce NMD activation ^58,59^. Higher TranslationAI scores at the downstream in-frame AUG were also associated with reduced predicted NMD-trigger probability, consistent with reports that translation reinitiation downstream of an early stop codon can reduce NMD sensitivity in specific transcript contexts ^60^. Together, these associations suggest that local translation context at both the early stop codon and downstream in-frame AUGs can contribute to the prediction of NMD-trigger probability.

The m6A-related features represent another category of novel predictors in predNMD. m6A is the most abundant internal modification in eukaryotic mRNA and has been implicated in multiple aspects of RNA metabolism, including splicing, export, stability, and translation ^61^. Recent studies have suggested potential connections between m6A modification and NMD. Notably, Ren et al., 2025 ^62^ demonstrated using direct RNA sequencing that NMD-targeted transcripts exhibit significantly higher m6A levels compared to their corresponding productive transcripts, both across the whole transcript and within the CDS region. Consistent with these observations, SHAP analysis showed that higher m6A density, both upstream of the early stop codon and across the whole transcript, was associated with increased NMD-trigger probability, supporting a potential functional relationship between m6A modification and NMD activity (Figure 1A).

The CAI difference feature captures local differences in codon optimality between the 25-codon window upstream of the early stop codon and the corresponding window upstream of the normal stop codon. SHAP analysis showed that higher CAI upstream of the early stop codon relative to the normal stop codon was associated with negative SHAP values, indicating lower predicted NMD-trigger probability (Figure 1A). This pattern points to a potential role for local codon optimality in NMD predictions, in line with broader evidence that codon usage can influence translation elongation and mRNA stability, although its specific relationship to “NMD efficiency” remains mechanistically unresolved ^63–65^

Spearman correlation analysis among the 20 selected features revealed interpretable clustering patterns based on hierarchical clustering of absolute correlation values (Figure 1B). There were four main clusters that stand out discussed from top to bottom in figure 1B. One cluster grouped mean gene expression, the two m6A density features, allele frequency, LOEUF, and phyloP score, which are features that reflect gene-level properties and evolutionary constraint rather than early stop codon-specific positional information. A second cluster contained the CDS position of the early stop codon and number of upstream exons, both of which capture how far the early stop codon is located from the start of translation and may be relevant to reinitiation potential. The third cluster grouped length of the exon containing early stop codon and distance to the end of the exon containing early stop codon, reflecting local exon architecture around the early stop codon. The fourth cluster contained features mostly related to the 3′ portion of the transcript: distance to 3′ UTR end, distance to the normal stop codon, the 50nt rule, number of downstream exons, and both TranslationAI scores. Notably, the two TranslationAI scores showed negative correlation with each other (ρ = −0.63), suggesting that early stop codons with strong termination signals tend to have weaker downstream reinitiation potential, and vice versa.

### ASE-derived NMD calls and predNMD predictions show position-dependent patterns across CDS regions

To visualize how NMD-associated allelic imbalance varies across coding sequence position, we examined the variant-level ASE measurements across GTEx, GREGOR, and GEUVADIS datasets, including variants later classified as indeterminate and excluded from model training (Figure 2A). TCGA was excluded from this ASE-level visualization because its labels were derived from transcript-level “NMD efficiency” rather than allele-specific expression. We summarized variants across three CDS regions: the first 400 nt downstream of the start codon, the 200 nt window flanking both sides of the last exon-exon junction, and the central CDS region, defined as the intervening region from 400 nt downstream of the start codon to 200 nt upstream of the last exon-exon junction. Because the length of this central region varies across transcripts, positions within the central CDS were rescaled to relative position units from 0 to 1.

**Figure 2.**
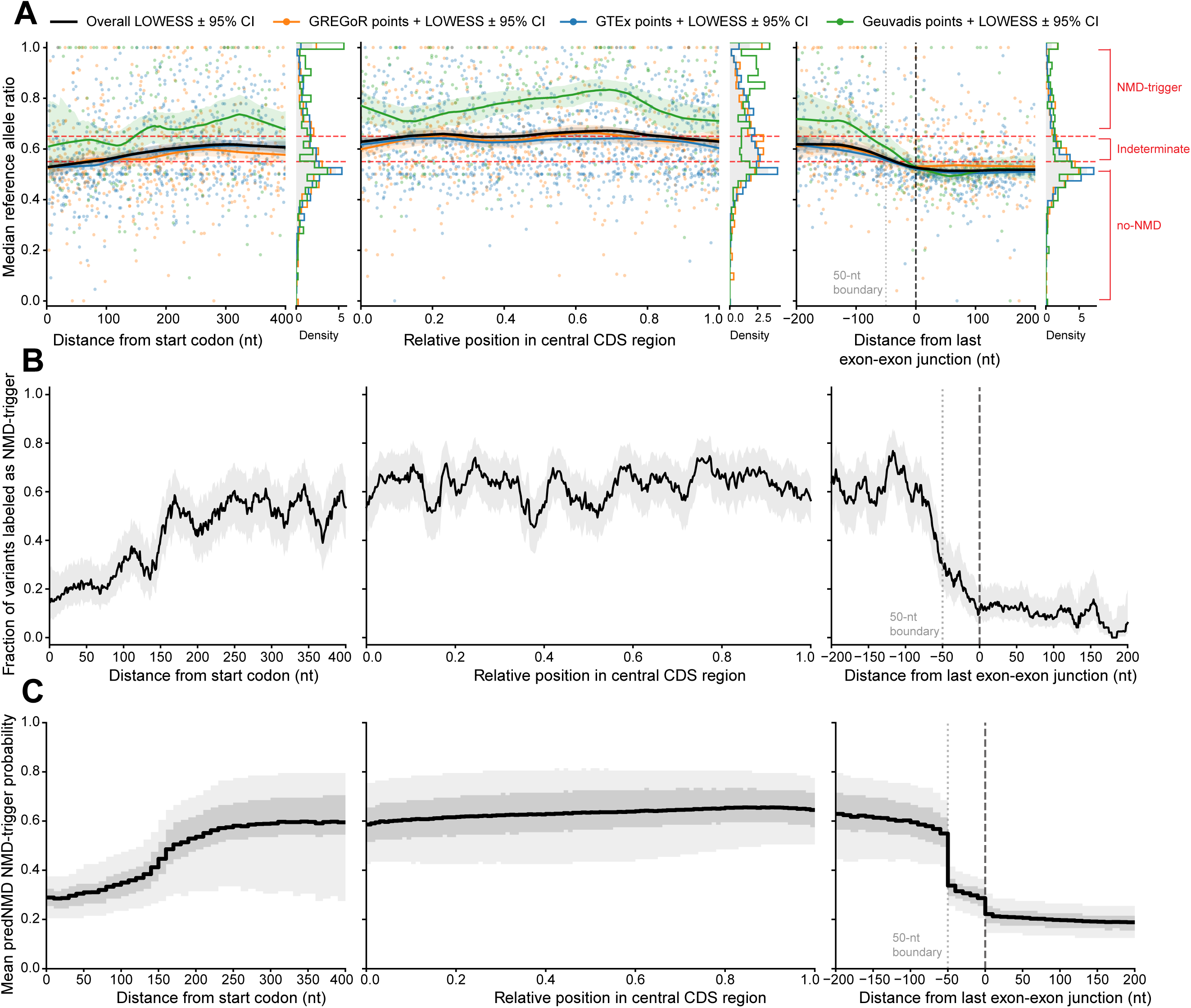
Positional patterns of allele-specific expression, training labels, and predNMD predictions across CDS regions. **(A)** Median reference allele ratio across the first 400 nt after the start codon, the central CDS region, and the ±200 nt window around the last exon-exon junction. Variants in the central CDS region were rescaled to relative CDS position from 0 to 1. Points show GREGoR, GTEx, and GEUVADIS variants colored by data source; TCGA is not shown because its labels were derived from transcript-level NMD efficiency rather than allele-specific expression. Lines show LOWESS trends with 95% confidence intervals. Marginal density plots show the distribution of median reference allele ratios within each region. Red dashed horizontal lines indicate the thresholds used to define no-NMD (<0.55) and NMD-trigger (>0.65) labels, with variants between these thresholds treated as indeterminate and excluded from the training dataset. **(B)** Fraction of binarized training variants labeled as NMD-trigger across the same CDS regions, including variants from GREGoR, GTEx, GEUVADIS, and TCGA. Variants were summarized using sliding windows of 25 nt in the start-codon and last-exon-exon-junction regions, and 0.04 in relative-position units in the central CDS region. Gray shading indicates the Wilson 95% confidence interval for the binomial fraction in each window. **(C)** Mean predNMD NMD-trigger probability across approximately 14 million precomputed nonsense variants in the human genome. Predictions were summarized using 10-nt bins in the start-codon and last-exon-exon-junction regions, and 0.01 in relative-position units in the central CDS region. The black step line shows the mean predicted probability within each bin; light gray shading indicates the 10th–90th percentile range, and dark gray shading indicates the 30th–70th percentile range. Across all panels, the gray dotted vertical line marks the 50-nt boundary upstream of the last exon-exon junction, and the black dashed vertical line marks the last exon-exon junction.

For each variant, the reference allele ratio was calculated as the number of RNA-seq reads supporting the reference allele divided by the total number of reads supporting the reference and alternate alleles. Because the same variant could be observed in multiple individuals or samples, we used the median reference allele ratio across samples for each variant. Under this allele orientation, ratios close to 0.5 indicate approximately balanced allelic expression and little evidence of NMD, whereas higher ratios indicate allelic imbalance consistent with NMD-mediated degradation.

Across all three datasets, median reference allele ratios were close to 0.5 within the first ∼100–150 nt downstream of the start codon, indicating approximately balanced allelic expression in this start-proximal region. The ratios then increased toward approximately 0.6 by ∼200 nt downstream of the start codon. (Figure 2A) This pattern is consistent with reports that early stop codons close to the start codon may show reduced NMD sensitivity, potentially through translation reinitiation at downstream AUG codons ^26,28^. Ratios remained elevated and approximately stable across the central CDS and then decreased toward 0.5 less than ∼50 nt upstream of the last exon-exon junction, consistent with the canonical 50nt rule for EJC-dependent NMD ^2,4^. All three ASE-based datasets showed similar qualitative trends despite differences in biological source and experimental details, although GEUVADIS showed a higher baseline ratio across all regions. The reason for this offset is unclear and may reflect dataset-specific differences in sample type, assay sensitivity, cohort composition, etc.

In the final predNMD training set, the fraction of variants labeled as NMD-trigger showed similar position-dependent patterns across the same CDS regions (Figure 2B). This indicates that the binarization step preserved the major positional trends observed in the underlying allele-specific expression measurements.

We next applied predNMD to 13,968,776 possible nonsense variants spanning the human genome and summarized predicted NMD-trigger probabilities across the same three CDS regions (Figure 2C). The genome-wide prediction landscape closely recapitulated the patterns observed in both the ASE measurements and the binarized training labels, including reduced NMD-trigger probability near the start codon and near the last exon-exon junction. Together, these analyses show that predNMD reproduces the major position-dependent patterns of NMD activity genome-wide, extending the trends observed in the training labels to the full space of possible nonsense variants.

### predNMD achieves robust cross-validated performance and generalizes to independent validation data

Following the chromosome-holdout validation strategy used by Teran et al., 2021 ^34^, we evaluated predNMD using leave-one-chromosome-out cross-validation (LOCO-CV) on 5,304 high-quality variants curated from GREGoR, GEUVADIS, GTEx, and TCGA. predNMD achieved a mean LOCO-CV AUC of 0.785 ± 0.011, approximately 0.08 higher than the LOCO-CV AUC reported by Teran et al. for their GTEx-trained LASSO regression model ^34^.

To evaluate generalization to held-out data, we assessed predNMD performance on the MMRF-TARGET dataset, an independent validation set comprising 319 variants not used during training. predNMD achieved an AUC of 0.78 on this independent dataset (Figure 3), demonstrating that the strong cross-validation performance translates to robust generalization. We compared predNMD against existing NMD prediction methods on the MMRF-TARGET validation set. NMDetective-A achieved AUC = 0.76, NMDetective-B achieved AUC = 0.72, and the Ensembl VEP NMD prediction achieved AUC = 0.69 (Figure 3). The improvement of predNMD over the 50nt rule was substantial: predNMD gained 0.13 AUC points over the 50nt rule, nearly matching the 0.15 AUC-point gain of the 50nt rule over random classification. In terms of AUC above the no-skill baseline, predNMD therefore captured approximately 1.9-fold more discriminative signal than the canonical 50nt rule alone.

**Figure 3.**
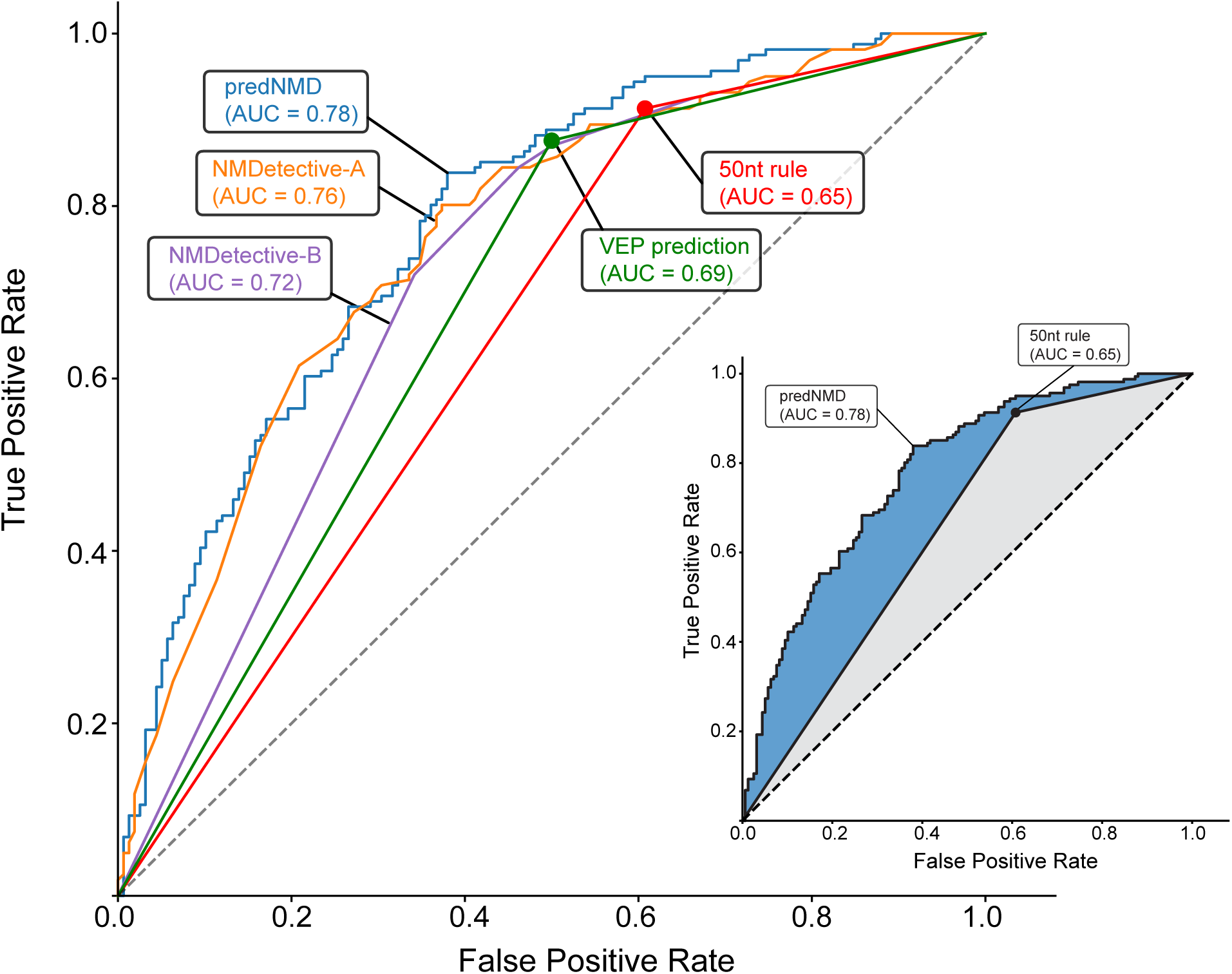
Performance comparison of NMD prediction methods on an independent validation set. Receiver operating characteristic (ROC) curves comparing predNMD against existing approaches, including NMDetective-A, NMDetective-B, VEP prediction, and the canonical 50nt rule, evaluated on the MMRF-TARGET dataset. The dashed diagonal line represents random classification, and area under the curve (AUC) values are shown for each method. Because the 50nt rule and VEP prediction produce binary classifications rather than continuous scores, they are shown as piecewise-linear ROC curves passing through their corresponding operating points. The inset highlights the direct comparison between predNMD and the 50nt rule. In the inset, the blue shaded region denotes the ROC area captured by predNMD above the 50nt rule, while the gray shaded region denotes the remaining ROC space above random classification but below the 50nt rule. Together, the shaded regions illustrate the relative discriminatory gain of predNMD over the canonical 50nt rule.

To assess whether the 20 selected features provided sufficient predictive information, we also compared performance between the model trained on all 166 initial features and the model trained on the 20 Boruta-selected features (Supplementary Figure 2). The 20-feature model achieved AUC = 0.78 compared to AUC = 0.76 for the 166-feature model on the MMRF-TARGET validation set. This result indicates that feature selection not only reduced model complexity but marginally improved generalization performance, likely by eliminating noisy or redundant features that could contribute to overfitting.

### predNMD predictions correlate with experimental measurements of transcript abundance

To evaluate whether predNMD’s predicted NMD-trigger probabilities reflect experimentally observed transcript abundance consequences, we tested the model in two saturation genome editing datasets where RNA readouts are available.

In the BRCA1 MAVE dataset ^43^, which contains 130 stop-gained mutations with RNA score measurements, predNMD’s NMD-trigger probability predictions showed strong correlation with reduced RNA scores (Pearson r = −0.60, p = 7.47×10⁻¹⁴; Spearman ρ = −0.64, p = 2.60×10⁻¹⁶) (Figure 4A). Mutations with high predNMD probability (>0.6) were generally associated with RNA scores below −1, corresponding to greater than 50% reduction in transcript abundance. Mutations with low predNMD probability (<0.4) were generally associated with RNA scores above −1, indicating less than 50% reduction. However, several mutations within 50 nt of the last exon-exon junction showed unexpectedly low RNA scores despite low predicted NMD-trigger probabilities; these mutations are examined further below. Overall, predNMD broadly distinguished mutations with stronger versus weaker RNA abundance reduction in the BRCA1 MAVE dataset.

**Figure 4.**
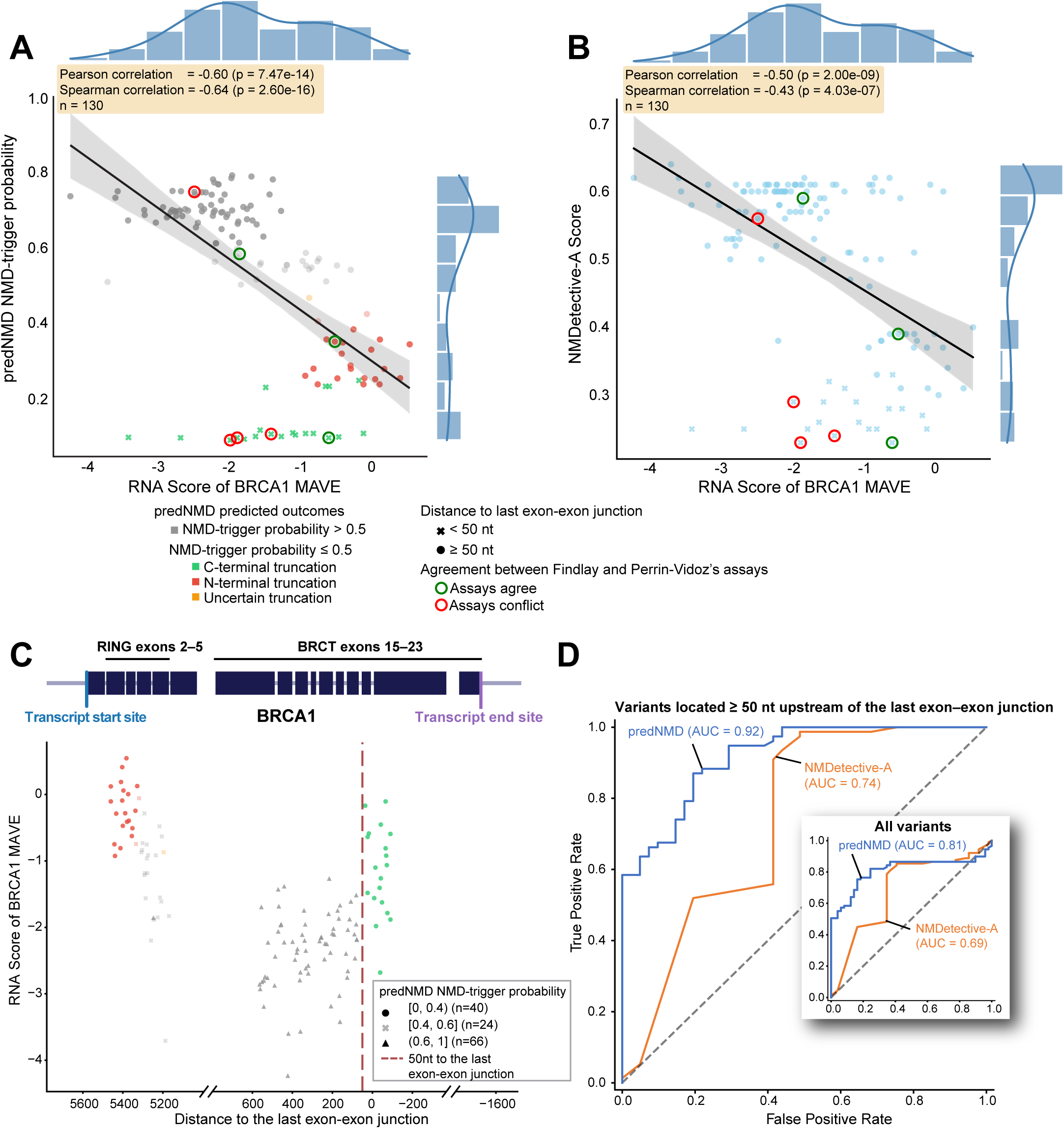
Correlation between NMD prediction score and RNA Score in the independent BRCA1 MAVE dataset. **(A)** Correlation between RNA Score and predNMD-predicted NMD-trigger probability across stop-gained variants (n = 130). Each point represents a variant and is colored by predNMD inferred truncation mechanism. A linear regression line with 95% confidence interval is shown. Pearson and Spearman correlation statistics are shown. Marginal histograms depict the distributions of RNA Score (top) and NMD-trigger probability (right). **(B)** Correlation between RNA Scores and NMDetective-A scores for the same set of variants (n = 130). Each point represents a variant, with the linear regression line and 95% confidence interval overlaid. Pearson and Spearman correlation statistics are shown. Marginal distributions of RNA Score (top) and NMDetective-A score (right) are displayed. Circled points denote variants where the assay outcome agrees (green) or disagrees (red) with the experimentally determined NMD status reported by Perrin-Vidoz et al., 2002. **(C)** RNA Score as a function of the distance from the early stop codon to the last exon–exon junction. Points are colored by truncation mechanism inferred by predNMD and shaped according to predNMD-predicted NMD-trigger probability bins. The dashed vertical line denotes the canonical 50nt rule boundary. **(D)** ROC curves evaluating predNMD and NMDetective-A as classifiers of NMD-triggering variants, using binarized RNA Scores (< −1: NMD-triggered; ≥ −1: no-NMD) as ground truth. The main plot shows performance across variants with early stop codons located ≥ 50 nt upstream of the last exon–exon junction. The inset shows performance across variants with early stop codons located ≥ 50 nt upstream of the last exon–exon junction. The inset shows performance on all 130 variants.

Mutations with intermediate predNMD probabilities (0.4–0.6) showed RNA scores distributed around −1, reflecting prediction uncertainty (Figure 4A). These mutations were predominantly located near the CDS start but downstream of the most start-proximal no-NMD mutations (Figure 4C). Notably, RNA scores showed a local decrease near a putative downstream AUG site, consistent with evidence that start-proximal NMD outcomes may depend on the feasibility of translation reinitiation, including the availability and spacing of downstream reinitiation sites ^66,67^. The distribution of RNA scores for these intermediate-probability mutations suggests that predNMD captures prediction uncertainty rather than forcing confident but potentially incorrect classifications.

Examination of mutations predicted to not trigger NMD revealed biologically coherent patterns (Figure 4C). We binarized predNMD outputs using an NMD-trigger probability threshold of 0.5, classifying mutations with predicted probability ≥ 0.5 as NMD-triggering and mutations with predicted probability < 0.5 as not triggering NMD. Among the 130 stop-gain BRCA1 mutations, predNMD classified 87 as NMD-triggering and 43 as not triggering NMD. Among the no-NMD mutations, 22 were predicted to produce N-terminal truncated protein, 20 were predicted to produce C-terminally truncated protein, and 1 showed uncertain mechanism assignment.

Unexpectedly, in the MAVE experiment, mutations located less than 50 nt upstream of the last exon-exon junction displayed variable RNA scores despite being predicted to not trigger NMD by the 50 nt rule. Some mutations very close to the normal stop codon exhibited surprisingly low RNA scores indicating approximately 90% reduction in mRNA abundance, despite NMD being highly unlikely at these positions according to all known mechanisms. To investigate whether this unexpected pattern reflects genuine biology or an experimental artifact of the MAVE assay, we compared MAVE RNA scores with the NMD status of several BRCA1 nonsense mutations experimentally determined by Perrin-Vidoz et al., 2002 ^68^ using cycloheximide treatment and quantitative RT-PCR. For some mutations, the two assays agreed on NMD status (green circles, Figure 4A,B), while others showed conflicting results (red circles, Figure 4A,B). Notably, the majority of mutations with conflicting labels between the two assays were located <50 nt upstream of the last exon-exon junction — precisely the subset exhibiting the unexpectedly spread-out RNA scores in the MAVE data. This concentration of inter-assay disagreements among last-exon mutations suggests that the surprisingly low MAVE RNA scores observed at these positions are more likely attributable to experimental artifacts of the MAVE assay rather than bona fide NMD, as the orthogonal cycloheximide-based assay by Perrin-Vidoz et al. did not confirm NMD-level transcript reduction for these mutations.

For comparison, we applied NMDetective-A scores to the same MAVE dataset since NMDetective-A achieved the second-best performance in our benchmark analysis on the MMRF-TARGET validation set. However, NMDetective-A scores showed substantially weaker correlation with RNA scores (Pearson r = −0.50, p = 2.00×10⁻⁹; Spearman ρ = −0.43, p = 4.03×10⁻⁷) (Figure 4B), and the separation between predicted NMD-triggering and no-NMD mutations was much less compelling compared to predNMD.

To quantify the classification performance of predNMD and NMDetective-A, we constructed ROC curves using binarized RNA scores as truth labels (RNA score < −1: NMD-triggering; ≥ −1: no-NMD) (Figure 4D). Because mutations located within 50 nt of the last exon-exon junction showed evidence of potential experimental artifacts in the MAVE assay, as discussed above, the main ROC analysis focused on mutations with early stop codons located ≥ 50 nt upstream of the last exon-exon junction, where the MAVE RNA scores are more likely to reflect genuine NMD status. In this subset, predNMD achieved an AUC of 0.92, outperforming NMDetective-A (AUC = 0.74) (Figure 4D). When evaluated across all 130 mutations, including the mutations near the last exon-exon junction with unexpectedly low MAVE RNA scores, predNMD maintained strong performance (AUC = 0.81) compared to NMDetective-A (AUC = 0.69) (Figure 4D), though the lower AUCs for both methods likely reflect noise introduced by unreliable truth labels among the last-exon mutations.

In the BARD1 MAVE dataset (Woo et al., 2025), which contains 318 stop-gained mutations with RNA score measurements, predNMD predictions again correlated well with experimental RNA scores and achieved high AUC (Pearson r = −0.57, p = 5.84×10⁻²⁹; AUC = 0.84). NMDetective-A showed substantially weaker correlation and lower AUC (Pearson r = −0.30, p = 4.30×10⁻⁸; AUC=0.70), further demonstrating predNMD’s improved predictive accuracy (Supplementary figure 3).

### predNMD shows stronger concordance with expert-curated clinical evidence than the 50nt rule

Accurate NMD prediction is central to applying PVS1, the strongest ACMG/AMP pathogenic evidence code, which current guidelines infer from the 50nt rule alone. Under the ACMG/AMP PVS1 framework, stop-gain variants predicted to trigger NMD may receive full PVS1 (i.e., with very strong evidence for pathogenicity) when loss of function is an established disease mechanism for the gene and the variant occurs in a biologically relevant isoform ^25^. In contrast, stop-gain variants predicted not to trigger NMD may receive reduced PVS1 strength or no PVS1 depending on the expected protein consequence. ClinGen Variant Curation Expert Panels (VCEPs), however, sometimes diverge from the standard ACMG/AMP decision tree and decline to apply full PVS1 for stop-gain variants that the 50nt rule predicts to trigger NMD, instead invoking gene-specific biological and clinical evidence. Such cases let us assess whether predNMD better reflects deliberate expert curation than the 50nt rule. We therefore examined predNMD predictions for stop-gain variants curated in the ClinGen Evidence Repository that were predicted to trigger NMD by the 50nt rule yet were not assigned the full PVS1 evidence code.

We focused on clinically informative cases in which the canonical 50nt rule predicted NMD triggering, but ClinGen VCEP curation did not assign full PVS1. After excluding variants for which the absence of full PVS1 was explained by factors unrelated to NMD prediction, such as lack of evidence that loss of function is the relevant disease mechanism, use of a clinically irrelevant isoform, or an explicit VCEP statement that the variant was still predicted to trigger NMD, eight informative variants remained. Among these cases, predNMD agreed with the VCEP interpretation for six variants by predicting no NMD triggering, whereas the 50nt rule predicted NMD triggering for all six. Thus, in the subset where the 50nt rule and VCEP interpretation disagreed for NMD-relevant reasons, predNMD aligned with expert curation for the majority of the variants.

Table 3 summarizes all curated variants from the ClinGen Evidence Repository, eight of which were informative, meaning the 50nt rule predicted NMD would be triggered but the VCEP did not assign full PVS1 for NMD-relevant reasons. For six of these eight, predNMD agreed with the VCEP while the 50nt rule did not; two occurred in APC: NM_000038.6(APC):c.70C>T (p.Arg24Ter) and NM_000038.6(APC):c.32dup (p.Gln12AlafsTer3). The InSiGHT Hereditary Colorectal Cancer/Polyposis VCEP developed an APC-specific PVS1 framework based on APC clinical evidence rather than the generic 50nt rule. Under this framework, PVS1 applies only to premature-truncation variants between codons 49 and 2645; variants upstream of codon 49 are considered not applicable for PVS1. This boundary was supported by extensive APC variant datasets, including 10,212 APC germline variants in ClinVar and 1,867 unique APC variants among 5,628 entries in the InSiGHT APC locus-specific database ^69^, together with literature review, expert elicitation, and pilot classification of 58 variants ^70^. The VCEP concluded that the absence of convincingly pathogenic loss-of-function variants upstream of codon 49 supports the interpretation that early stops in this region are unlikely to behave as canonical NMD-mediated loss-of-function alleles ^70^. Consistent with this VCEP model, predNMD predicted that both APC variants in the evidence repository upstream-of-codon 49 do not trigger NMD and predNMD inferred N-terminal truncation. The other four variants had ClinGen evidence noting that early stop codons within the first 100 nucleotides may not trigger NMD because of translation reinitiation, again consistent with predNMD prediction of N-terminal truncation. These results indicate that predNMD captures biological reasoning used in expert curation for a subset of variants where the 50nt rule alone would assign NMD-trigger status.

**Table 3.**
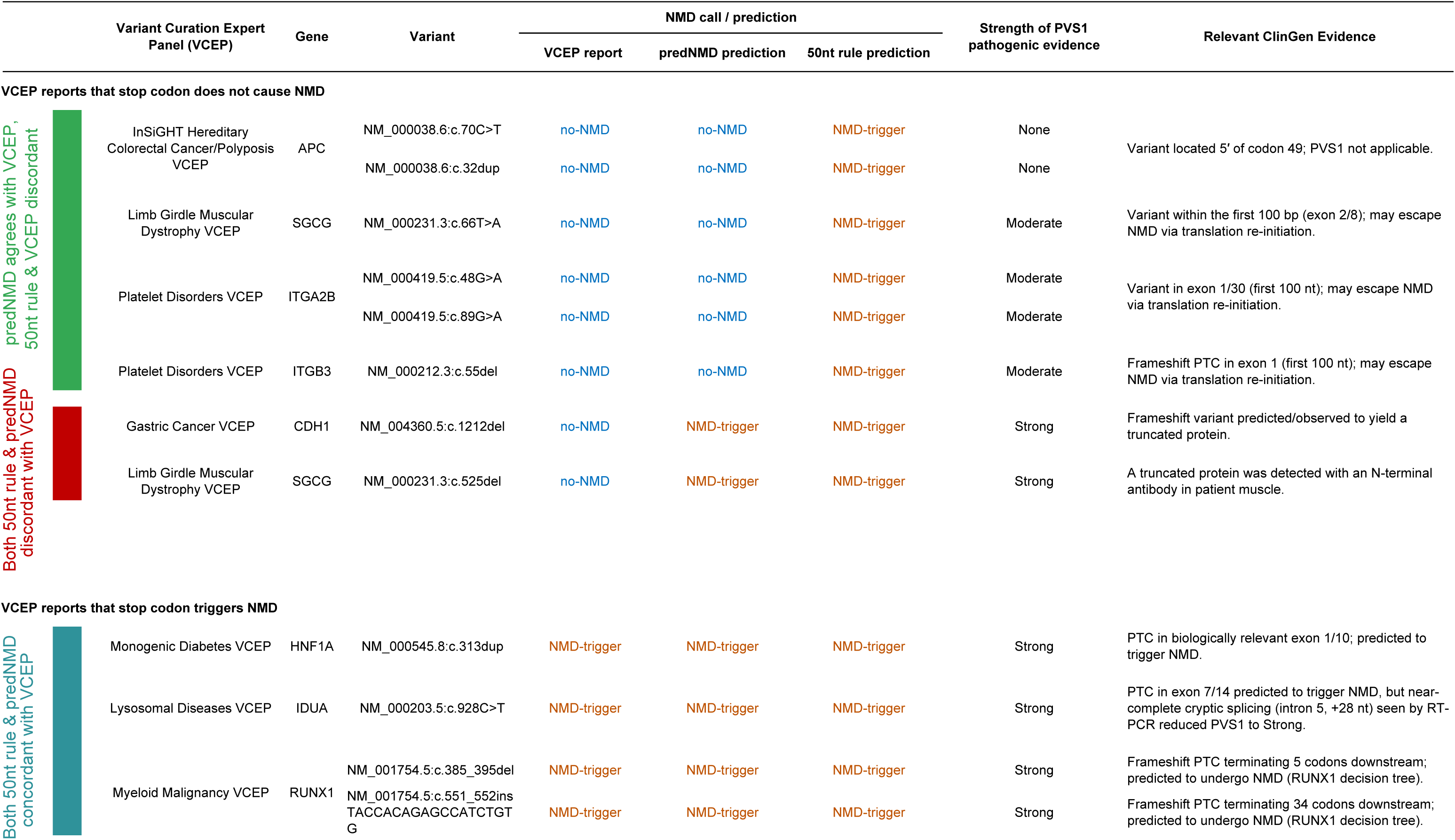
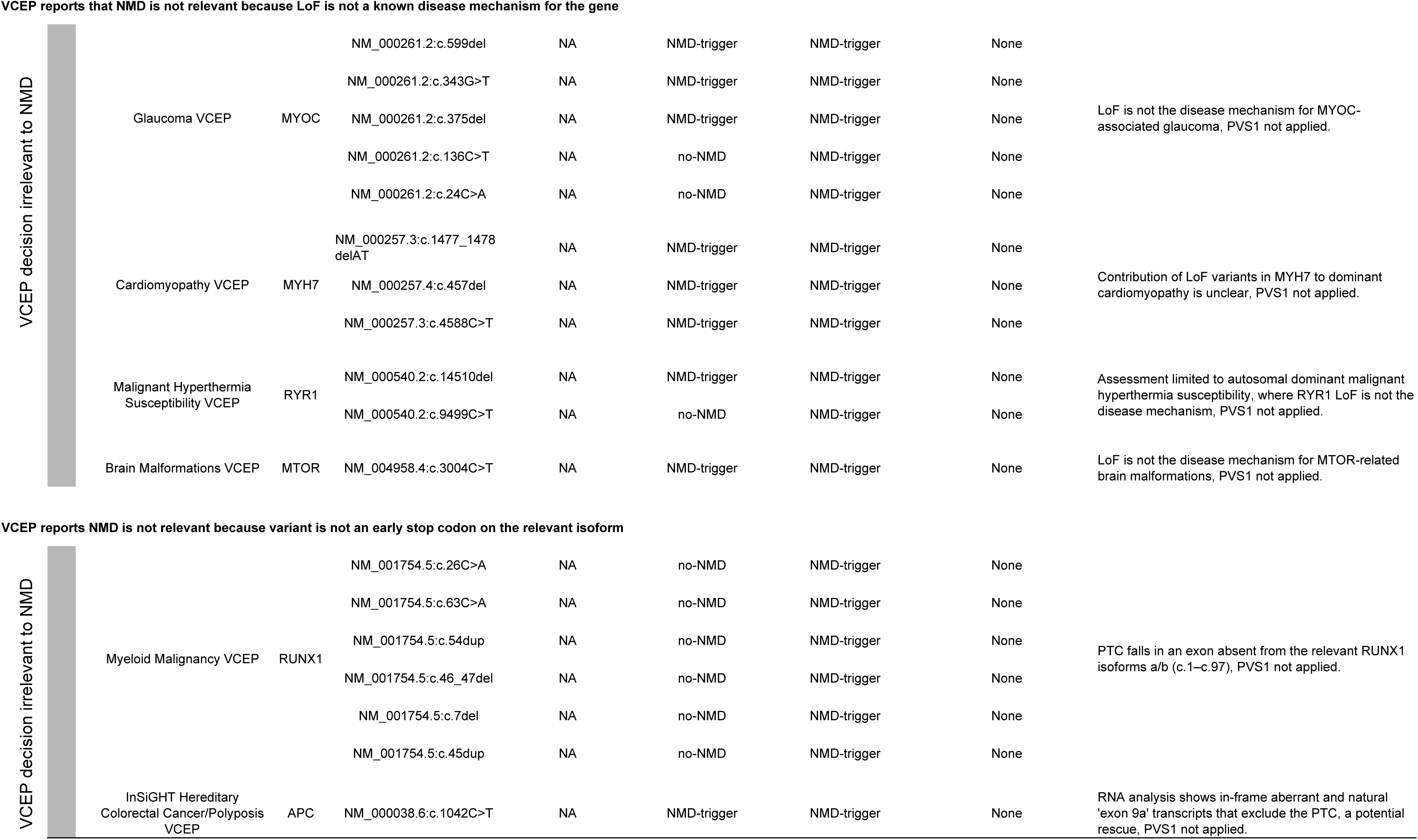
Stop gain variants >50nt upstream of last exon-exon junction in ClinGen Evidence Repository for which the VCEP did not assign full PVS1 Very Strong.

The remaining two informative variants, NM_004360.5(CDH1):c.1212del and NM_000231.3(SGCG):c.525del, were predicted to trigger NMD by both predNMD and the 50nt rule, yet the VCEP concluded that each yields a truncated protein and therefore assigned PVS1 at Strong rather than full Very Strong. For the SGCG variant, the Limb Girdle Muscular Dystrophy VCEP cited direct functional evidence: a truncated protein detected with an N-terminal antibody in patient muscle, indicating that the allele produces stable protein. For the CDH1 variant, the Gastric Cancer VCEP likewise interpreted the variant as producing a truncated protein, although the Evidence Repository does not record the basis for this conclusion.

Together, this analysis shows that predNMD improves agreement with expert curation in clinically informative discordant cases. In the subset where the 50nt rule and VCEP interpretation disagreed for NMD-relevant reasons, predNMD agreed with the VCEP interpretation for six of eight variants, supporting its value as a more clinically consistent model of NMD prediction beyond the canonical 50nt rule.

## Discussion

In this study, we developed predNMD, a method for predicting nonsense-mediated mRNA decay outcomes trained on large datasets and incorporating novel predictive features. predNMD achieved robust cross-validated performance and outperformed existing NMD prediction methods on independent validation data. Beyond binary NMD prediction, predNMD provides interpretable predictions of the likely protein truncation mechanism when NMD does not occur, enabling more nuanced clinical variant interpretation. Validation against MAVE datasets and expert-curated clinical evidence demonstrated that predNMD captures biologically meaningful signals and can support variant curation decisions in cases where the canonical 50nt rule alone is insufficient.

### Recommendation: incorporating predNMD into clinical workflows for stop-gain variant classification

A key motivation for developing predNMD is to improve assessment of stop-gain variants within the ACMG/AMP PVS1 frameworks ^25^ and the forthcoming ACMG/AMP/CAP/ClinGen Sequence Variant Classification (SVC) v4.0 clinical variant classification framework Loss of Function decision tree ^71^. In the current ACMG/AMP guidelines, the PVS1 decision tree for nonsense and frameshift variants includes a critical branch point at which variants are evaluated for whether they are “predicted to undergo NMD” ^25^. This determination currently relies on the 50nt rule, which as demonstrated by our results and those of others, does not capture the full complexity of NMD regulation.

We therefore propose that predNMD be incorporated into clinical stop-gain variant classification as a refined NMD-prediction module at the existing “predicted to undergo NMD” decision node in the PVS1 framework and SVC v4.0 successors. This requires no change to the overall structure of the PVS1 decision tree. Instead, predNMD would be a “drop-in” replacement for the 50nt rule as the primary method for estimating whether an annotated early termination codon is likely to trigger NMD. Variants with high predNMD NMD-trigger probability would proceed through the rest of the existing PVS1 branch for predicted NMD-triggering variants, including gene-disease mechanism and biologically relevant transcript assessment required by ACMG/AMP and VCEP-specific specifications. Variants with low predNMD NMD-trigger probability would proceed through the existing branch for variants not predicted to undergo NMD, where PVS1 strength may be reduced depending on the expected truncated protein consequence.

Although data are limited to calibrate predNMD, this recommendation is supported by the magnitude of predNMD’s improvement over the 50nt rule. On the independent MMRF-TARGET validation set, predNMD achieved an AUC of 0.78, compared with 0.65 for the 50nt rule (Figure 3). The 0.13 AUC improvement of predNMD over the 50nt rule nearly matched the 0.15 AUC improvement of the 50nt rule over random classification, indicating that predNMD captures almost as much additional discriminatory signal beyond the 50nt rule as the 50nt rule captures beyond random prediction. In ClinGen-curated stop-gain variants not assigned full PVS1, predNMD matched expert-panel judgments in most cases where the 50nt rule predicted NMD would be triggered but the panels concluded, from additional evidence, that these variants were unlikely to trigger NMD. The rich gene-specific evidence that informs VCEP decisions to depart from the 50nt rule, such as the large APC locus-specific database that underpinned the InSiGHT VCEP’s codon 49 boundary, is available to only a handful of VCEPs. Most expert panels lack comparable data and therefore default to the 50nt rule, motivating accurate predictors that improve classification consistency across the many genes where such bespoke evidence is unavailable. Together, these results suggest that replacing the 50nt rule with predNMD at the NMD decision step would make PVS1 interpretation more consistent with both observed NMD biology and expert clinical reasoning.

To support direct clinical and research use, we provide precomputed predNMD predictions for for every possible stop-gain single-nucleotide variant, totaling 13,968,776 for Ensembl GRCh38 (release 104) and 12,714,784 for GRCh37 (release 87). These precomputed predictions can be accessed through the publicly available predNMD web interface at predNMD.org or queried in batch using indexed precomputed tables. For users querying a small number of variants, individual stop-gain variants can be entered directly into the web interface to retrieve predNMD predictions. For larger variant sets, we provide a lightweight Unix/Linux shell script with no additional software dependencies. Given the predNMD precompute file and a query VCF, the script extracts stop-gain variants from the VCF, looks them up in the predNMD precomputed index, and writes a tab-delimited output containing the stop-gain variants with predNMD predictions. The batch-query workflow processed a full genomic VCF in 30 seconds on one CPU core of a 2021 desktop PC with an Intel Core i7-10700 CPU at 2.90 GHz and 32 GB RAM. predNMD is released under an open-source license and is freely available for academic, clinical, and research use.

In addition to using precomputed predictions, users may wish to run the full software locally when analyzing custom annotations, alternative transcript sets, or future reference genome and gene annotation releases. For this purpose, predNMD is distributed as both an installable Python package and a Docker container, ensuring compatibility across diverse computational environments and reproducibility of results.

Together, these features make predNMD straightforward to adopt on two complementary levels. Conceptually, predNMD serves as a drop-in refinement at the existing “predicted to undergo NMD” decision node of the PVS1 framework and the forthcoming SVC v4.0 standard, replacing the 50nt rule, without altering the surrounding decision tree. Practically, predNMD can be run by labs and clinicians with minimal infrastructure, whether through precomputed predictions, the installable Python package, the Docker container, or the predNMD.org web interface.

### m6A modification emerges as a candidate modulator of NMD outcome

Our feature selection analysis identified several novel characteristics of NMD outcomes, including m6A density, TranslationAI-derived scores, and local sequence features. The relationship between m6A modification and NMD warrants further investigation, as our results suggest both promoting and inhibiting effects that may depend on the spatial distribution of m6A marks relative to the early stop codon.

Previous studies have established that m6A is not uniformly distributed across transcripts but instead exhibits characteristic spatial patterns. Uzonyi et al., 2023 ^72^ demonstrated that m6A is largely depleted within 100-200 nt of either side of splice sites (defined as exclusion zone) and enriched in the middle regions of exons longer than 400 nt. This observation provides a potential mechanistic explanation for the long-standing but unexplained finding that early stop codons located in unusually long exons or far from exon boundaries are less likely to trigger NMD ^33,34^. We hypothesize that early stop codons positioned in the m6A-enriched central regions of long exons may escape NMD, possibly through steric competition of m6A reader proteins (such as YTHDF and YTHDC family members) with EJC deposition or UPF1 recruitment, though the precise mechanism remains to be determined.

Several previous studies have identified specific points of contact between m6A and RNA surveillance, including NMD. Li et al., 2019 ^73^ showed in glioblastoma that METTL3-dependent m6A modulates the NMD sensitivity of splicing-factor transcripts and thereby influences alternative-splicing isoform switches. He et al., 2024 ^74^ reported in plants that m6A-modified PepMV RNA is recognized by ECT2-family readers that interact with the NMD-related factors UPF3 and SMG7, and that the antiviral effect of m6A depends on these NMD-related factors. Tan et al., 2021 ^75^ showed that processed pseudogene transcripts, which lack introns and are therefore poor substrates for canonical EJC-dependent NMD, have elevated m6A levels, rapidly evolved m6A motifs, and undergo m6A-promoted cytosolic RNA degradation, consistent with a complementary surveillance route for NMD-resistant RNAs. Ren et al., 2025 ^62^ further found that highly methylated misprocessed isoforms, including NMD-class isoforms, are enriched for m6A especially within coding regions, further implicating m6A in surveillance of aberrant isoforms.

To explore this hypothesis, we analyzed the distribution of m⁶A sites within the exon containing the early stop codon in our training data, stratified by NMD outcome. We compared m6A counts between NMD-triggering and no-NMD variants located > 50 nt upstream of the last exon-exon junction (Supplementary Figure 4). No-NMD variants showed a trend toward higher m6A enrichment within the exon containing early stop codon (mean: 5 ± 17) compared to NMD-triggering variants (mean: 3 ± 11), consistent with a model in which m⁶A modifications in the vicinity of the early stop codon may protect transcripts from NMD. However, this difference did not reach statistical significance (two-sided Mann–Whitney U test; p = 0.075), potentially due to the limited sample size available in our training data and the high variance in m6A counts within both groups. Nonetheless, the observed trend is consistent with prior findings linking m⁶A modifications to NMD evasion, and larger sample sizes or enhanced experimental methods may be needed to provide sufficient statistical power to detect this effect.

These observations present a seeming paradox: local m6A enrichment near the early stop codon appears to promote NMD escape, while our model also shows that higher overall m6A density across the transcript is associated with increased NMD probability, consistent with Ren et al., 2025 ^62^. One possible explanation is that global and local m6A marks play distinct roles, global m6A enrichment may mark transcripts for general surveillance and turnover, while local m6A accumulation near early stop codon may interfere with NMD-specific machinery assembly. Alternatively, the correlation between global m6A levels and NMD may not be causal but rather reflect shared features of NMD-targeted transcript isoforms. Resolving the precise mechanisms by which m6A influences NMD will require targeted experimental studies that manipulate m6A deposition at specific transcript positions.

### Candidate features not retained after Boruta feature selection

We conducted a feature selection process that retained only a small fraction of the candidate features we developed (retaining 20 of 166; Table 2, Supplementary Figure 1). Several candidate features that were not selected were motivated by prior evidence implicating them in NMD or translation termination. These exclusions are informative because they help distinguish molecular signals with broad, genome-wide predictive value from signals that may be context-specific, redundant with retained features, or poorly represented by the available feature definitions. Many excluded features were redundant variants of retained ones, and their exclusion is unsurprising. For example, there were alternative versions of m6A density and CAI difference, each computed over a different region or window size. Some excluded categories are more notable, however, given their established links to NMD or translation termination, and we discuss these below.

We first considered features related to stop-codon readthrough, a known mechanism by which transcripts can escape NMD. These comprised traditional readthrough-context features extensively studied as determinants of readthrough efficiency ^76,77^. These candidate features were: the stop-codon identity (UGA, UAG, or UAA), the nucleotide at the −3 position and whether it is a U, and the nucleotides at the +4, +5, and +9 positions, where the stop codon occupies positions +1 to +3 and downstream nucleotides are numbered accordingly. All readthrough-related features were eliminated during selection. It is possible that the TranslationAI termination score at the early stop codon, which was retained, may already capture termination efficiency in a way that subsumes the contribution of these individual nucleotide-context features. Efficient readthrough is also relatively rare at most early stop codons, which may further limit the genome-wide predictive value of these features.

We also developed RNA-binding protein (RBP) binding candidate features, comprising both known RBP motifs and de novo motifs identified within the region from the early stop codon to the 3′ UTR end. These were motivated by evidence that RBPs binding near the termination codon can modulate NMD susceptibility; for example, PTBP1 and hnRNP L bound just downstream of a stop codon reported to block UPF1 accumulation on the 3′ UTR and protect long-3′ UTR transcripts from decay ^78,79^. None of the RBP motif features were chosen by the Boruta selection, suggesting that motif-based occupancy predictions did not add signal beyond the retained features. We are unsure why they were not recognized as having significant discriminative information, though we note that matching of predicted motif is an imperfect proxy for actual in vivo RBP binding.

We included the GC content of the 200 nt immediately downstream of the early stop codon, motivated by reports that GC-rich sequences in the 3′ UTR promote UPF1-dependent mRNA decay ^80^. Although whole-transcript GC content was retained, this local downstream GC feature was not.

Finally, we included the length of the 5′ UTR, as 5′ UTR architecture has been implicated in translation initiation and efficiency ^81^. This feature was not selected; unlike 3′ UTR length, 5′ UTR length is a coarse transcript-level descriptor with no established direct role in NMD, and its exclusion is consistent with prior NMD predictors, which have not adopted it as a feature.

### Binary NMD classification reflects clinical convention

predNMD is designed as a binary classifier that predicts whether a variant triggers NMD rather than the magnitude of protein reduction. This design choice reflects the structure of current clinical guidelines: in the ACMG/AMP framework, NMD is treated as a binary decision point in the PVS1 decision tree, where variants either are or are not expected to undergo NMD, with corresponding implications for evidence code assignment ^25^. However, an important biological distinction exists between the probability that a variant triggers NMD and the degree of mRNA and protein reduction when NMD does occur. “NMD efficiency” can vary substantially, with some transcripts experiencing near-complete degradation while others show only partial reduction. The relationship between transcript level and protein output is complicated. NMD limits protein accumulation through mechanisms beyond transcript degradation, including translational repression, so that transcripts escaping degradation may persist yet yield little protein ^82^. Conversely, downstream homeostatic responses can buffer the functional consequences of an NMD event. Many genes tune their own expression through autoregulatory feedback. At the transcriptional level a gene product can feed back to modulate its own transcription rate ^83^. Post-transcriptionally, this can occur by several means, including coupling alternative splicing to NMD ^15,22,84^. Separately, degradation of the mutant transcript can itself trigger transcriptional adaptation, a form of genetic compensation that upregulates paralogous genes independently of protein loss ^85,86^. predNMD predicts the molecular triggering event and cannot anticipate these systemic responses. Future clinical interpretation will therefore benefit from models that predict not only whether NMD is triggered but also the expected magnitude of protein reduction, enabling more refined assessments of residual protein function. Developing such models will require systematic protein-level measurements across many variants, which remain technically challenging to obtain at scale. Moreover, translating continuous predictions of protein reduction into clinical practice would require corresponding updates to the variant interpretation framework, which currently treats NMD as a binary decision point. Defining where a protein-reduction threshold should be drawn, and how graded efficiency estimates map onto discrete evidence strength levels, would in turn require detailed, gene- and protein-specific knowledge of how much residual protein is functionally sufficient.

### Potential limitations and future directions

Stop-gain variants make up only a small fraction of the genetic variation carried in any individual genome, yet they are substantially enriched among variants of established clinical significance and account for a disproportionate share of known pathogenic variants ^87^. Improving the interpretation of these variants therefore has an impact on clinical genomics far larger than their genomic frequency would suggest, and predNMD represents a step toward that goal. Fully determining the impact of stop-gain variants, however, will depend on data resources that do not yet exist at sufficient scale. Each marks a concrete direction in which coordinated, community-wide data generation could enable substantially more sophisticated models for NMD prediction, and enhancement of predNMD.

Most fundamentally, large-scale cohorts rarely measure NMD directly, and therefore predNMD was trained on indirect proxies for NMD activity. ASE infers NMD from allelic imbalance in transcript abundance, but such imbalance can also arise from allele-specific transcription, epigenetic modification, or altered splicing ^88–90^, and ASE is further subject to noise from sequencing depth, mapping bias, and biological variation. Transcript-level “NMD efficiency” estimates, such as those used in the TCGA-derived training data from Lindeboom et al. ^33^, share this indirectness and add a further caveat. Derived from tumors, they may not reflect germline or normal-tissue NMD, as copy-number alterations, aneuploidy, altered NMD-factor expression, and clonal heterogeneity can all distort transcript abundance relative to the constitutional context in which most clinically interpreted variants act. Lindeboom et al. ^33^ applied extensive filtering and denoising procedures to reduce technical and biological confounders in these estimates, but differences in transcript abundance remain non-specific to NMD. In addition, because these proxies reflect steady-state transcript abundance and NMD is frequently incomplete, the presence of residual mutant transcript does not guarantee that the variant does not trigger NMD; such transcript may be only partially degraded or not yet cleared, so variants that do trigger NMD may be mislabeled as non-triggering. To limit label ambiguity, we used a double-threshold scheme rather than the single threshold of previous methods, labeling variants only when their values fell clearly below or above two separate cutoffs and excluding intermediate, indeterminate variants. Combining four independent resources (GTEx, TCGA, GEUVADIS, and GREGoR) increases variant number and diversity and reduces single-source bias, although these datasets are not perfectly calibrated to one another, as the GEUVADIS baseline offset in Figure 2A illustrates. Residual label noise is therefore inevitable and likely bounds achievable model performance.

A related limitation stems from these training data: because predNMD is trained on binary labels derived from heterogeneous, noisy proxies rather than direct NMD measurements, its NMD-trigger probabilities are model-estimated values rather than fully calibrated biological probabilities. Label noise tends to compress these predicted probabilities toward intermediate values, pulling them away from 0 and 1 even for variants whose true outcome is near-certain. Consistent with this, the binned mean predNMD NMD-trigger probability across CDS regions closely follows the fraction of training variants labeled as NMD-trigger across the same positional regions (Figure 2B,C). The resulting scores should therefore be interpreted as relative NMD-trigger likelihoods rather than calibrated measurements of biological “NMD efficiency”. Future work using more reliable, less noisy experimental datasets could enable explicit probability calibration, so that predNMD probabilities correspond more closely to observed NMD-trigger frequencies across variant and transcript contexts.

Experimental MAVE datasets, which we used as complementary validation for predNMD, are also imperfect proxies for NMD. These assays are gene-specific, rely on engineered variant libraries, and may be biased toward particular transcript regions depending on assay design, editable sequence contexts, and the distribution of assayed variants. In addition to NMD-mediated transcript depletion, RNA scores may also reflect assay-specific effects and other biological events impacting transcript abundance as described above ^43,44^. For example, in the BRCA1 MAVE dataset, several variants located <50 nt of the last exon-exon junction showed unexpectedly low RNA scores even though their position is not expected to trigger NMD under any known rule and an independent assay ^68^ confirmed no NMD at these positions, suggesting that even direct experimental RNA readouts may contain context-specific or assay-specific artifacts. A further practical limitation is the scarcity of usable MAVE data: relatively few MAVEs have been performed in a genomic rather than cDNA context, and fewer still include RNA or protein readouts informative for NMD, which constrains both the quantity and the genomic representativeness of MAVE-based validation.

The consequence of a stop-gain variant can also depend on which transcript isoform harbors it: genes often express multiple isoforms, and a variant that triggers NMD on one isoform may not on another because of differences in exon structure or 3′ UTR length. Unambiguously determining which isoforms are expressed in relevant tissues, however, often requires long-read sequencing data that are not yet available at sufficient scale to resolve full-length transcript structures genome-wide. In the absence of such data, predNMD evaluates each variant in the context of a single representative transcript, the canonical isoform of its gene, rather than the full set of isoforms the gene expresses. Integration of long-read data to model isoform-specific NMD outcomes represents an important direction for future model development.

Finally, the degree to which NMD varies across biological contexts remains debated. On one hand, Zetoune et al. 2008 reported tissue-specific differences in NMD efficiency in mice, with mutant transcript levels varying more than two-fold across tissues ^91^; inter-individual variation in “NMD efficiency” has been proposed to contribute to phenotypic variability among patients carrying the same pathogenic variant ^92^; and NMD activity changes during development, including dynamic regulation during fly embryogenesis, mammalian brain development, myogenesis, and B-cell maturation ^93^. On the other hand, Teran et al., 2021 ^34^ found NMD effects to be highly concordant across tissues and individuals in GTEx data, suggesting that binary NMD triggering may be largely consistent even when the magnitude of transcript reduction varies. These conflicting findings leave the degree of context-dependence unresolved, and modeling context-specific NMD variation remains a challenge for future work.

While these considerations define the current boundaries of our approach, predNMD represents a significant advance over existing NMD prediction methods in terms of prediction accuracy. Furthermore, the ability to infer truncation mechanisms for variants not triggering NMD adds clinically relevant information for distinguishing haploinsufficiency from potential dominant-negative effects. We anticipate that predNMD will be useful for improving application of the PVS1 criterion in clinical variant interpretation and for prioritizing variants for experimental follow-up. The predNMD software, Docker container, and precomputed predictions for all possible single nucleotide stop-gained variants are freely available to facilitate broad adoption and support streamlined integration into research and clinical genomics workflows.

## Acknowledgements

This project was initiated in the ClinGen Sequence Variant Interpretation Working Group, and we thank its members and those of the ClinGen Computational Working Group. We thank all the Brenner lab members for their feedback and suggestions. We thank Dr. Rik G. H. Lindeboom for sharing the processed TCGA dataset. We thank Dr. Sujatha Jagannathan for her suggestions. This study was supported by Tata Consultancy Services and by NIH grants U24 HG007346, P01 AI138962.

## Generative AI usage

Generative AI tools were used to assist with code development and text refinement; the authors reviewed and validated all output and take full responsibility for the content.

## Note

We have been made aware that there is related work to build a modern NMD predictor led by Dr. Zeynep Coban Akdemir in collaboration with Dr. Sujatha Jagannathan.

## Data and code availability

No new primary sequencing data were generated in this study. The predNMD source code is available at https://github.com/BrennerLab/predNMD. A prebuilt Docker image containing the predNMD runtime environment and required dependencies is available at https://hub.docker.com/r/brennerlab/prednmd. Precomputed predNMD predictions and the batch-query shell script are both available through the predNMD web interface at https://predNMD.org.

---

**Supplementary Figure 1.**
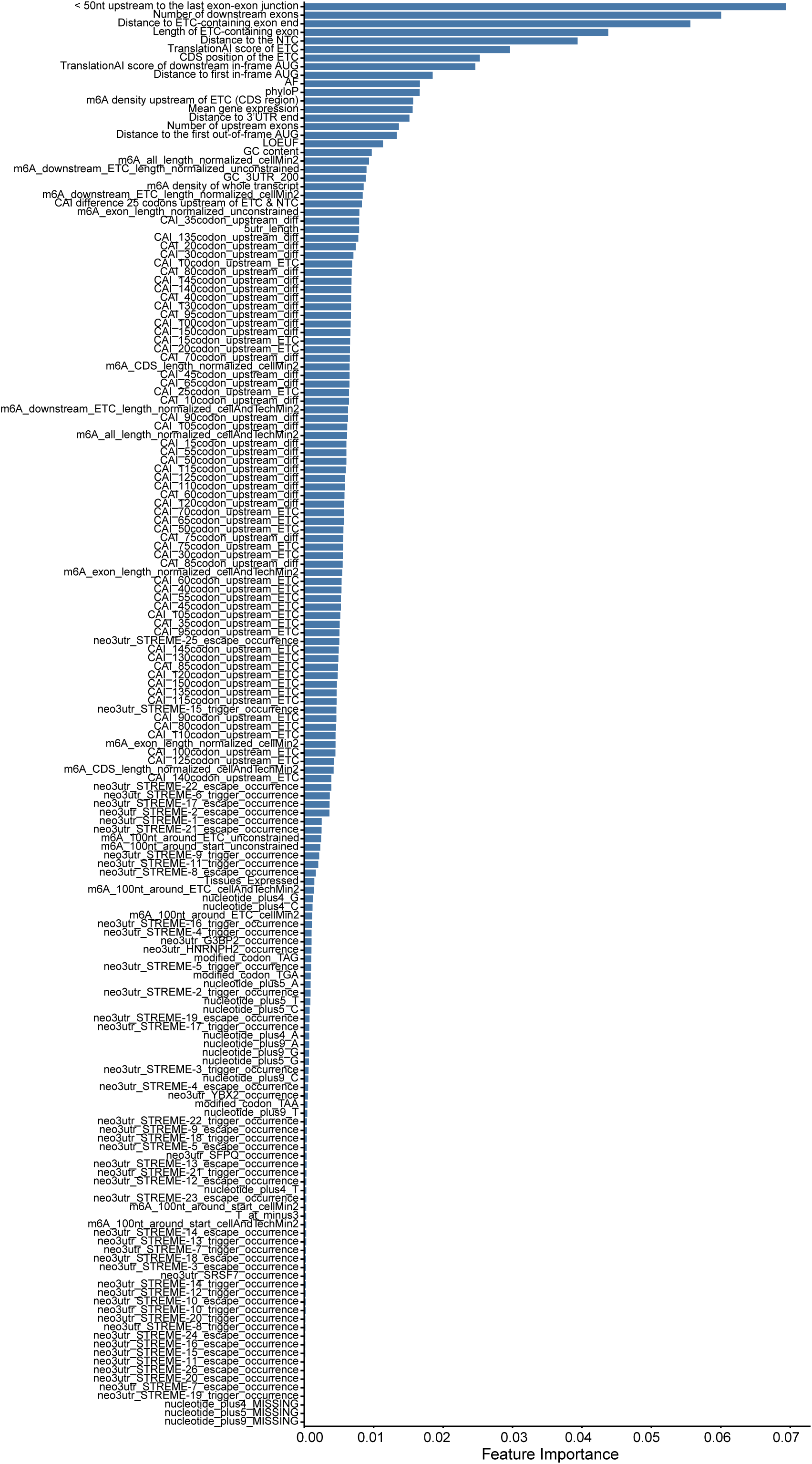
Mean Decrease Impurity importance scores of the full 166 features initially included in predNMD before feature selection.

**Supplementary Figure 2.**
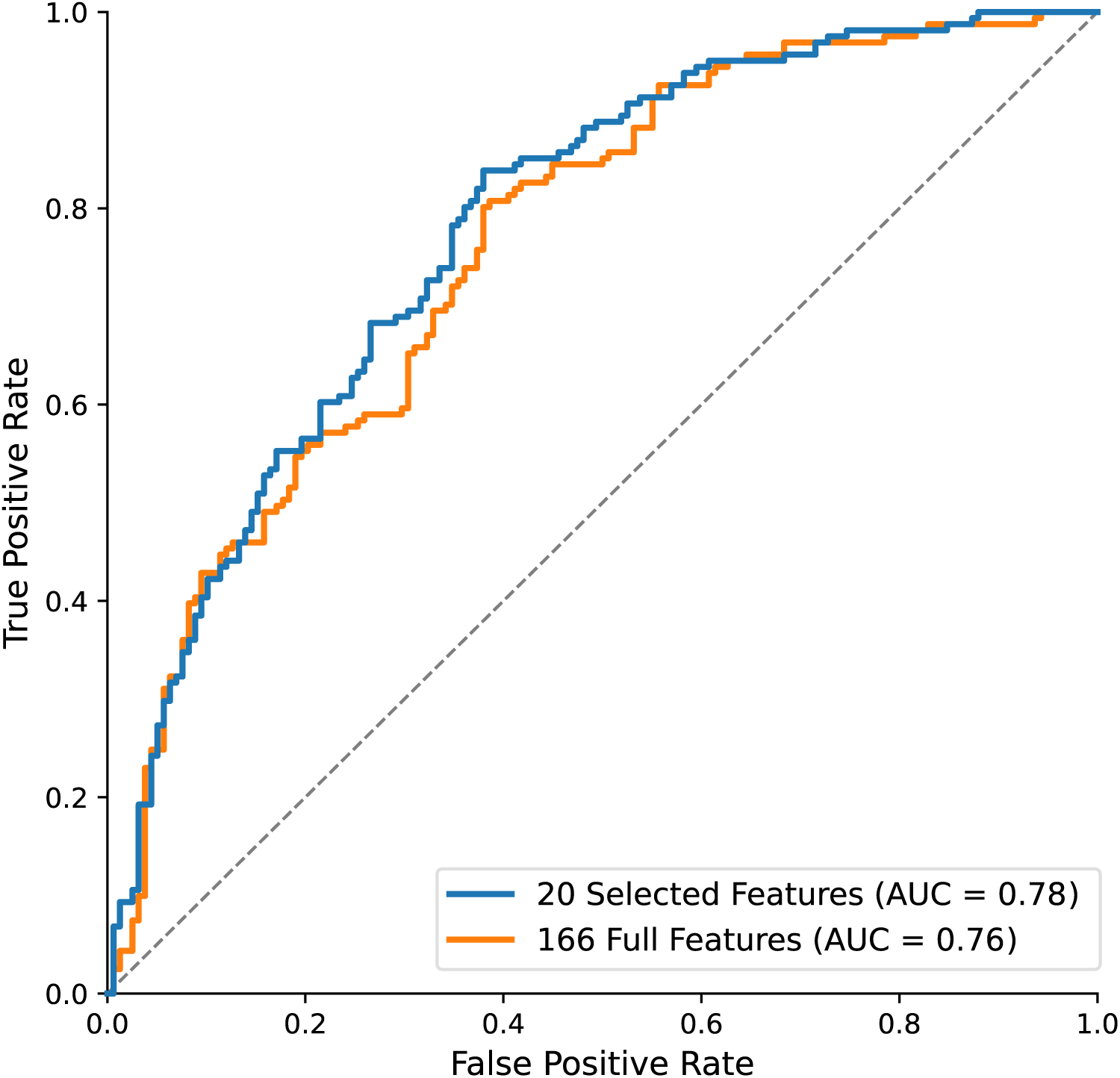
ROC curves comparison between Random Forest model trained with the full 166 features and model trained with the 20 selected features on the independent validation set. Dashed diagonal line represents random classification. Area under the curve (AUC) values are shown for each model.

**Supplementary Figure 3.**
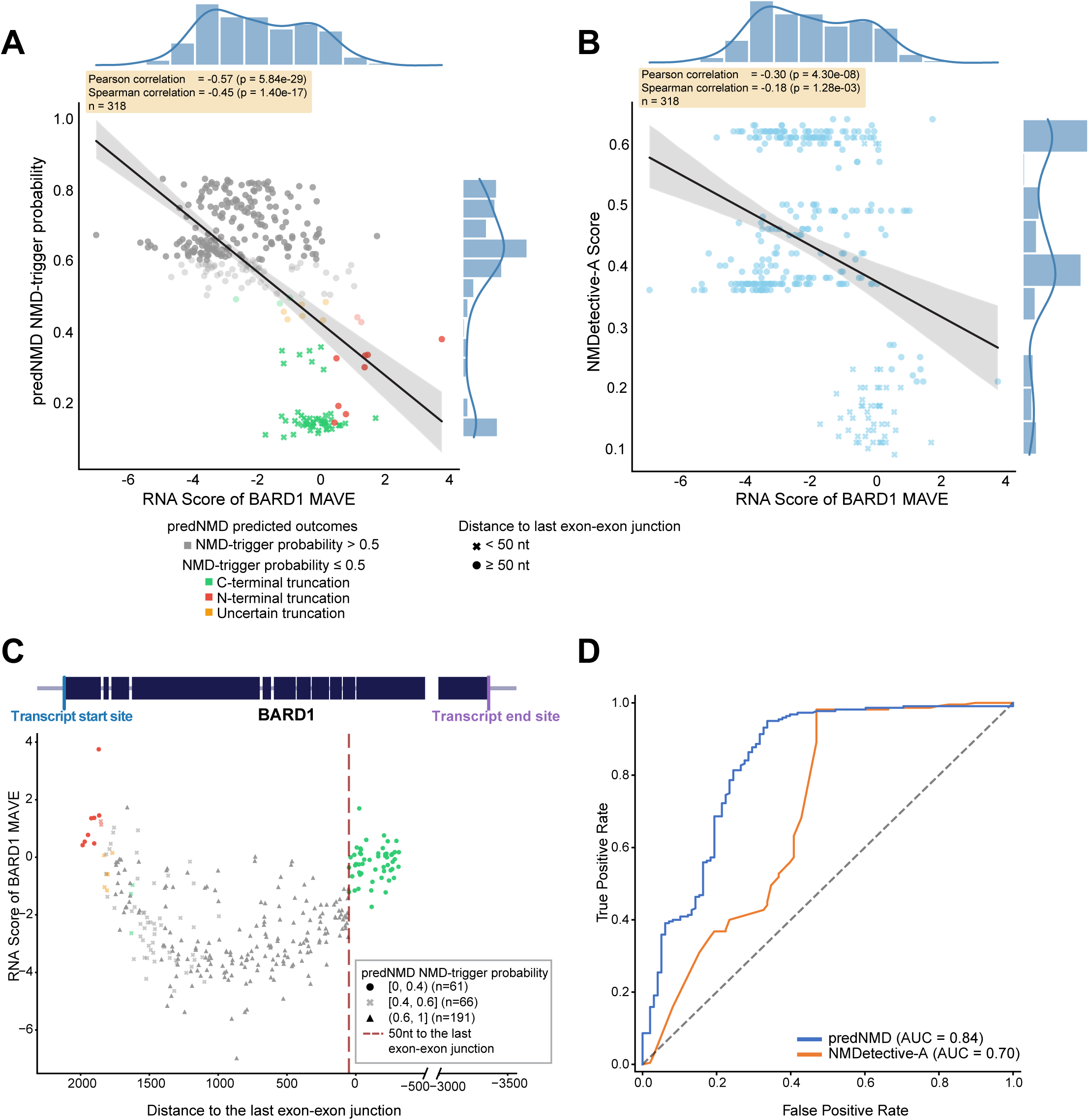
Correlation between NMD prediction score and RNA Score in the independent BARD1 MAVE dataset. **(A)** Correlation between RNA Score and predNMD-predicted NMD-trigger probability across stop-gained variants (n = 318). Each point represents a variant and is colored by predNMD inferred truncation mechanism. A linear regression line with 95% confidence interval is shown. Pearson correlation coefficients and corresponding p-values are indicated. Marginal histograms depict the distributions of RNA Score (top) and NMD-trigger probability (right). **(B)** Correlation between RNA Scores and NMDetective-A scores for the same set of variants (n = 318). Each point represents a variant, with the linear regression line and 95% confidence interval overlaid. Pearson correlation statistics are shown. Marginal distributions of RNA Score (top) and NMDetective-A score (right) are displayed. **(C)** RNA Score as a function of the distance from the early stop codon to the last exon–exon junction. Points are colored by truncation mechanism inferred by predNMD and shaped according to predNMD NMD-trigger probability bins. The dashed vertical line denotes the canonical 50nt rule boundary. **(D)** ROC curves evaluating predNMD and NMDetective-A as classifiers of NMD-triggering variants, using binarized RNA Scores (< −1: NMD-triggered; ≥ −1: no-NMD) as ground truth. The main plot shows performance across variants with early stop codons located ≥ 50 nt upstream of the last exon–exon junction. The inset shows performance on all 318 variants.

**Supplementary Figure 4.**
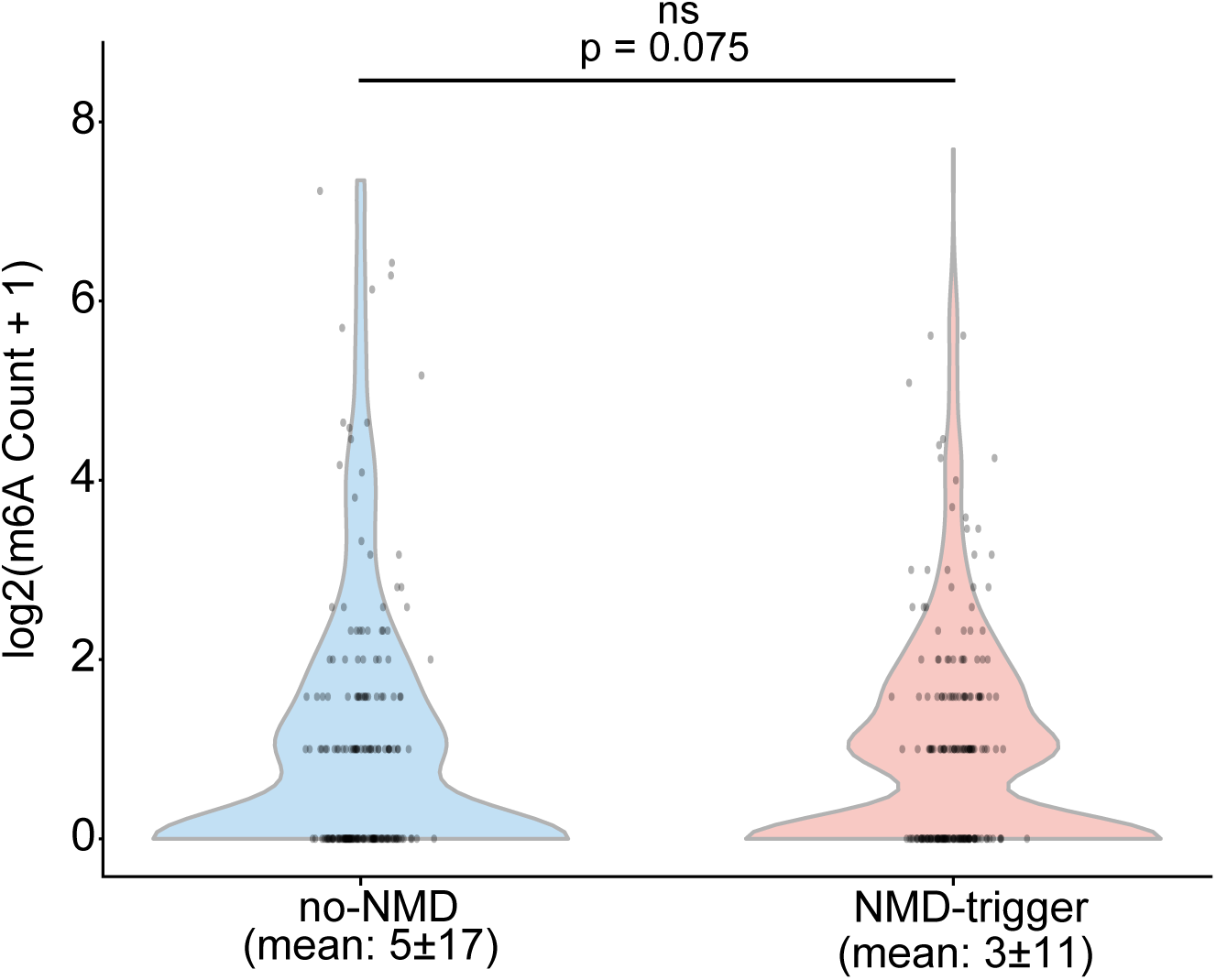
Differential distribution of m⁶A sites within the exon containing early stop codon between NMD-triggering and no-NMD variants. Distributions of m6A counts (log2[count + 1]) are shown for NMD-triggering and no-NMD variants located > 50nt upstream of the last exon-exon junction. Violin plots display the full distributions with individual variants overlaid. NMD-triggering variants show reduced m6A enrichment within the exon containing early stop codon compared with no-NMD variants, but the difference is not statistically significant (two-sided Mann–Whitney U test; p = 0.075).

## References

1. Leeds, P., Peltz, S.W., Jacobson, A., and Culbertson, M.R. (1991). The product of the yeast UPF1 gene is required for rapid turnover of mRNAs containing a premature translational termination codon. Genes Dev 5, 2303–2314.

2. Nagy, E., and Maquat, L.E. (1998). A rule for termination-codon position within intron-containing genes: when nonsense affects RNA abundance. Trends Biochem Sci 23, 198–199.

3. Mitchell, P., and Tollervey, D. (2001). mRNA turnover. Curr Opin Cell Biol 13, 320–325.

4. Le Hir, H., Moore, M.J., and Maquat, L.E. (2000). Pre-mRNA splicing alters mRNP composition: evidence for stable association of proteins at exon-exon junctions. Genes Dev 14, 1098–1108.

5. Ishigaki, Y., Li, X., Serin, G., and Maquat, L.E. (2001). Evidence for a pioneer round of mRNA translation: mRNAs subject to nonsense-mediated decay in mammalian cells are bound by CBP80 and CBP20. Cell 106, 607–617.

6. Lykke-Andersen, J., Shu, M.D., and Steitz, J.A. (2001). Communication of the position of exon-exon junctions to the mRNA surveillance machinery by the protein RNPS1. Science 293, 1836–1839.

7. Herskowitz, I. (1987). Functional inactivation of genes by dominant negative mutations. Nature 329, 219–222.

8. Kurosaki, T., Popp, M.W., and Maquat, L.E. (2019). Quality and quantity control of gene expression by nonsense-mediated mRNA decay. Nat Rev Mol Cell Biol 20, 406–420.

9. Holbrook, J.A., Neu-Yilik, G., Hentze, M.W., and Kulozik, A.E. (2004). Nonsense-mediated decay approaches the clinic. Nat Genet 36, 801–808.

10. Inoue, K., Khajavi, M., Ohyama, T., Hirabayashi, S.-I., Wilson, J., Reggin, J.D., Mancias, P., Butler, I.J., Wilkinson, M.F., Wegner, M., et al. (2004). Molecular mechanism for distinct neurological phenotypes conveyed by allelic truncating mutations. Nat Genet 36, 361–369.

11. Monaco, A.P., Bertelson, C.J., Liechti-Gallati, S., Moser, H., and Kunkel, L.M. (1988). An explanation for the phenotypic differences between patients bearing partial deletions of the DMD locus. Genomics 2, 90–95.

12. Chelly, J., Gilgenkrantz, H., Lambert, M., Hamard, G., Chafey, P., Récan, D., Katz, P., de la Chapelle, A., Koenig, M., and Ginjaar, I.B. (1990). Effect of dystrophin gene deletions on mRNA levels and processing in Duchenne and Becker muscular dystrophies. Cell 63, 1239–1248.

13. Kerr, T.P., Sewry, C.A., Robb, S.A., and Roberts, R.G. (2001). Long mutant dystrophins and variable phenotypes: evasion of nonsense-mediated decay? Hum Genet 109, 402–407.

14. Crawford, G.E., Faulkner, J.A., Crosbie, R.H., Campbell, K.P., Froehner, S.C., and Chamberlain, J.S. (2000). Assembly of the dystrophin-associated protein complex does not require the dystrophin COOH-terminal domain. J Cell Biol 150, 1399–1410.

15. Lewis, B.P., Green, R.E., and Brenner, S.E. (2003). Evidence for the widespread coupling of alternative splicing and nonsense-mediated mRNA decay in humans. Proc Natl Acad Sci U S A 100, 189–192.

16. Sureau, A., Gattoni, R., Dooghe, Y., Stévenin, J., and Soret, J. (2001). SC35 autoregulates its expression by promoting splicing events that destabilize its mRNAs. EMBO J 20, 1785–1796.

17. Desai, A., Hu, Z., French, C., Lloyd, J., and Brenner, S. (2020). Networks of splice factor regulation by unproductive splicing coupled with Nonsense mediated mRNA decay. bioRxiv. 10.1101/2020.05.20.107375.

18. Carvill, G.L., Engel, K.L., Ramamurthy, A., Cochran, J.N., Roovers, J., Stamberger, H., Lim, N., Schneider, A.L., Hollingsworth, G., Holder, D.H., et al. (2018). Aberrant Inclusion of a Poison Exon Causes Dravet Syndrome and Related SCN1A-Associated Genetic Epilepsies. Am J Hum Genet 103, 1022–1029.

19. Voskobiynyk, Y., Battu, G., Felker, S.A., Cochran, J.N., Newton, M.P., Lambert, L.J., Kesterson, R.A., Myers, R.M., Cooper, G.M., Roberson, E.D., et al. (2021). Aberrant regulation of a poison exon caused by a non-coding variant in a mouse model of Scn1a-associated epileptic encephalopathy. PLoS Genet 17, e1009195.

20. Lim, K.H., Han, Z., Jeon, H.Y., Kach, J., Jing, E., Weyn-Vanhentenryck, S., Downs, M., Corrionero, A., Oh, R., Scharner, J., et al. (2020). Antisense oligonucleotide modulation of non-productive alternative splicing upregulates gene expression. Nat Commun 11, 3501.

21. Leclair, N.K., Brugiolo, M., Park, S., Devoucoux, M., Urbanski, L., Angarola, B.L., Yurieva, M., and Anczuków, O. (2025). Antisense oligonucleotide-mediated TRA2β poison exon inclusion induces the expression of a lncRNA with anti-tumor effects. Nat Commun 16, 1670.

22. Lareau, L.F., Inada, M., Green, R.E., Wengrod, J.C., and Brenner, S.E. (2007). Unproductive splicing of SR genes associated with highly conserved and ultraconserved DNA elements. Nature 446, 926–929.

23. Lareau, L.F., and Brenner, S.E. (2015). Regulation of splicing factors by alternative splicing and NMD is conserved between kingdoms yet evolutionarily flexible. Mol Biol Evol 32, 1072–1079.

24. Richards, S., Aziz, N., Bale, S., Bick, D., Das, S., Gastier-Foster, J., Grody, W.W., Hegde, M., Lyon, E., Spector, E., et al. (2015). Standards and guidelines for the interpretation of sequence variants: a joint consensus recommendation of the American College of Medical Genetics and Genomics and the Association for Molecular Pathology. Genet Med 17, 405–424.

25. Abou Tayoun, A.N., Pesaran, T., DiStefano, M.T., Oza, A., Rehm, H.L., Biesecker, L.G., Harrison, S.M., and ClinGen Sequence Variant Interpretation Working Group (ClinGen SVI) (2018). Recommendations for interpreting the loss of function PVS1 ACMG/AMP variant criterion. Hum Mutat 39, 1517–1524.

26. Zhang, J., and Maquat, L.E. (1997). Evidence that translation reinitiation abrogates nonsense-mediated mRNA decay in mammalian cells. EMBO J 16, 826–833.

27. Hogg, J.R., and Goff, S.P. (2010). Upf1 senses 3’UTR length to potentiate mRNA decay. Cell 143, 379–389.

28. Lindeboom, R.G.H., Supek, F., and Lehner, B. (2016). The rules and impact of nonsense-mediated mRNA decay in human cancers. Nat Genet 48, 1112–1118.

29. Hurt, J.A., Robertson, A.D., and Burge, C.B. (2013). Global analyses of UPF1 binding and function reveal expanded scope of nonsense-mediated mRNA decay. Genome Res 23, 1636–1650.

30. Hoek, T.A., Khuperkar, D., Lindeboom, R.G.H., Sonneveld, S., Verhagen, B.M.P., Boersma, S., Vermeulen, M., and Tanenbaum, M.E. (2019). Single-Molecule Imaging Uncovers Rules Governing Nonsense-Mediated mRNA Decay. Mol Cell 75, 324–339.e11.

31. Kolakada, D., Campbell, A.E., Galvis, L.B., Li, Z., Lore, M., and Jagannathan, S. (2024). A system of reporters for comparative investigation of EJC-independent and EJC-enhanced nonsense-mediated mRNA decay. Nucleic Acids Res 52, e34–e34.

32. Rivas, M.A., Pirinen, M., Conrad, D.F., Lek, M., Tsang, E.K., Karczewski, K.J., Maller, J.B., Kukurba, K.R., DeLuca, D.S., Fromer, M., et al. (2015). Human genomics. Effect of predicted protein-truncating genetic variants on the human transcriptome. Science 348, 666–669.

33. Lindeboom, R.G.H., Vermeulen, M., Lehner, B., and Supek, F. (2019). The impact of nonsense-mediated mRNA decay on genetic disease, gene editing and cancer immunotherapy. Nat Genet 51, 1645–1651.

34. Teran, N.A., Nachun, D.C., Eulalio, T., Ferraro, N.M., Smail, C., Rivas, M.A., and Montgomery, S.B. (2021). Nonsense-mediated decay is highly stable across individuals and tissues. Am J Hum Genet 108, 1401–1408.

35. Kim, Y.-G., Kang, H., Lee, B., Jang, H.-J., Park, J.-H., Ha, C., Park, H., and Kim, J.-W. (2024). A spectrum of nonsense-mediated mRNA decay efficiency along the degree of mutational constraint. Commun Biol 7, 1461.

36. GTEx Consortium (2013). The Genotype-Tissue Expression (GTEx) project. Nat Genet 45, 580–585.

37. Cancer Genome Atlas Research Network, Weinstein, J.N., Collisson, E.A., Mills, G.B., Shaw, K.R.M., Ozenberger, B.A., Ellrott, K., Shmulevich, I., Sander, C., and Stuart, J.M. (2013). The Cancer Genome Atlas Pan-Cancer analysis project. Nat Genet 45, 1113–1120.

38. Lappalainen, T., Sammeth, M., Friedländer, M.R., ‘t Hoen, P.A.C., Monlong, J., Rivas, M.A., Gonzàlez-Porta, M., Kurbatova, N., Griebel, T., Ferreira, P.G., et al. (2013). Transcriptome and genome sequencing uncovers functional variation in humans. Nature 501, 506–511.

39. Dawood, M., Heavner, B., Wheeler, M.M., Ungar, R.A., LoTempio, J., Wiel, L., Berger, S., Bernstein, J.A., Chong, J.X., Délot, E.C., et al. (2025). GREGoR: accelerating genomics for rare diseases. Nature 647, 331–342.

40. McLaren, W., Gil, L., Hunt, S.E., Riat, H.S., Ritchie, G.R.S., Thormann, A., Flicek, P., and Cunningham, F. (2016). The Ensembl Variant Effect Predictor. Genome Biol 17, 122.

41. Castel, S.E., Levy-Moonshine, A., Mohammadi, P., Banks, E., and Lappalainen, T. (2015). Tools and best practices for data processing in allelic expression analysis. Genome Biol 16, 195.

42. Rubin, A.F., Stone, J., Bianchi, A.H., Capodanno, B.J., Da, E.Y., Dias, M., Esposito, D., Frazer, J., Fu, Y., Grindstaff, S.B., et al. (2025). MaveDB 2024: a curated community database with over seven million variant effects from multiplexed functional assays. Genome Biol 26, 13.

43. Findlay, G.M., Daza, R.M., Martin, B., Zhang, M.D., Leith, A.P., Gasperini, M., Janizek, J.D., Huang, X., Starita, L.M., and Shendure, J. (2018). Accurate classification of BRCA1 variants with saturation genome editing. Nature 562, 217–222.

44. Woo, I., Casadei, S., Snyder, M.W., Smith, N.T., Best, S., Tejura, M., Gupta, P., McEwen, A.E., Hamm, A., Dawood, M., et al. (2025). Saturation genome editing of BARD1 resolves VUS and provides insight into BRCA1-BARD1 tumor suppression.

45. Pedregosa, F., Varoquaux, G., Gramfort, A., Michel, V., Thirion, B., Grisel, O., Blondel, M., Müller, A., Nothman, J., Louppe, G., et al. (2012). Scikit-learn: Machine Learning in Python.

46. Kursa, M.B., Jankowski, A., and Rudnicki, W.R. (2010). Boruta – A System for Feature Selection. Fundamenta Informaticae.

47. Liang, Z., Ye, H., Ma, J., Wei, Z., Wang, Y., Zhang, Y., Huang, D., Song, B., Meng, J., Rigden, D.J., et al. (2023). m6A-Atlas v2.0: updated resources for unraveling the N6-methyladenosine (m6A) epitranscriptome among multiple species. Nucleic Acids Res 52, D194–D202.

48. Athey, J., Alexaki, A., Osipova, E., Rostovtsev, A., Santana-Quintero, L.V., Katneni, U., Simonyan, V., and Kimchi-Sarfaty, C. (2017). A new and updated resource for codon usage tables. BMC Bioinformatics 18, 391.

49. Sharp, P.M., and Li, W.H. (1987). The codon Adaptation Index--a measure of directional synonymous codon usage bias, and its potential applications. Nucleic Acids Res 15, 1281–1295.

50. Karczewski, K.J., Francioli, L.C., Tiao, G., Cummings, B.B., Alföldi, J., Wang, Q., Collins, R.L., Laricchia, K.M., Ganna, A., Birnbaum, D.P., et al. (2020). The mutational constraint spectrum quantified from variation in 141,456 humans. Nature 581, 434–443.

51. Pollard, K.S., Hubisz, M.J., Rosenbloom, K.R., and Siepel, A. (2010). Detection of nonneutral substitution rates on mammalian phylogenies. Genome Res 20, 110–121.

52. Fan, X., Chang, T., Chen, C., Hafner, M., and Wang, Z. (2025). Analysis of RNA translation with a deep learning architecture provides new insight into translation control. Nucleic Acids Res 53, gkaf277.

53. Lundberg, S., and Lee, S.-I. (2017). A Unified Approach to Interpreting Model Predictions.

54. Lundberg, S.M., Erion, G., Chen, H., DeGrave, A., Prutkin, J.M., Nair, B., Katz, R., Himmelfarb, J., Bansal, N., and Lee, S.-I. (2020). From local explanations to global understanding with explainable AI for trees. Nature Machine Intelligence 2, 56–67.

55. Bridle, J.S. (1990). Probabilistic interpretation of feedforward classification network outputs, with relationships to statistical pattern recognition. In Neurocomputing, (Berlin, Heidelberg: Springer Berlin Heidelberg), pp. 227–236.

56. Le Hir, H., Izaurralde, E., Maquat, L.E., and Moore, M.J. (2000). The spliceosome deposits multiple proteins 20-24 nucleotides upstream of mRNA exon-exon junctions. EMBO J 19, 6860–6869.

57. Kashima, I., Yamashita, A., Izumi, N., Kataoka, N., Morishita, R., Hoshino, S., Ohno, M., Dreyfuss, G., and Ohno, S. (2006). Binding of a novel SMG-1-Upf1-eRF1-eRF3 complex (SURF) to the exon junction complex triggers Upf1 phosphorylation and nonsense-mediated mRNA decay. Genes Dev 20, 355–367.

58. Annibaldis, G., Domanski, M., Dreos, R., Contu, L., Carl, S., Kläy, N., and Mühlemann, O. (2020). Readthrough of stop codons under limiting ABCE1 concentration involves frameshifting and inhibits nonsense-mediated mRNA decay. Nucleic Acids Res 48, 10259–10279.

59. Embree, C.M., Abu-Alhasan, R., and Singh, G. (2022). Features and factors that dictate if terminating ribosomes cause or counteract nonsense-mediated mRNA decay. J Biol Chem 298, 102592.

60. Neu-Yilik, G., Amthor, B., Gehring, N.H., Bahri, S., Paidassi, H., Hentze, M.W., and Kulozik, A.E. (2011). Mechanism of escape from nonsense-mediated mRNA decay of human beta-globin transcripts with nonsense mutations in the first exon. RNA 17, 843–854.

61. Roundtree, I.A., Evans, M.E., Pan, T., and He, C. (2017). Dynamic RNA Modifications in Gene Expression Regulation. Cell 169, 1187–1200.

62. Ren, Z., He, J., Huang, X., Gao, Y., Wei, C., Wu, Z., Guo, W., Wang, F., Zhao, Q., Sun, X., et al. (2025). Isoform characterization of m6A in single cells identifies its role in RNA surveillance. Nature Communications 16, 5828.

63. Presnyak, V., Alhusaini, N., Chen, Y.-H., Martin, S., Morris, N., Kline, N., Olson, S., Weinberg, D., Baker, K.E., Graveley, B.R., et al. (2015). Codon optimality is a major determinant of mRNA stability. Cell 160, 1111–1124.

64. Radhakrishnan, A., Chen, Y.-H., Martin, S., Alhusaini, N., Green, R., and Coller, J. (2016). The DEAD-Box Protein Dhh1p Couples mRNA Decay and Translation by Monitoring Codon Optimality. Cell 167, 122–132.e9.

65. Wu, Q., Medina, S.G., Kushawah, G., DeVore, M.L., Castellano, L.A., Hand, J.M., Wright, M., and Bazzini, A.A. (2019). Translation affects mRNA stability in a codon-dependent manner in human cells. Elife 8,.

66. Pereira, F.J.C., Teixeira, A., Kong, J., Barbosa, C., Silva, A.L., Marques-Ramos, A., Liebhaber, S.A., and Romão, L. (2015). Resistance of mRNAs with AUG-proximal nonsense mutations to nonsense-mediated decay reflects variables of mRNA structure and translational activity. Nucleic Acids Res 43, 6528–6544.

67. Cohen, S., Kramarski, L., Levi, S., Deshe, N., Ben David, O., and Arbely, E. (2019). Nonsense mutation-dependent reinitiation of translation in mammalian cells. Nucleic Acids Res 47, 6330–6338.

68. Perrin-Vidoz, L., Sinilnikova, O.M., Stoppa-Lyonnet, D., Lenoir, G.M., and Mazoyer, S. (2002). The nonsense-mediated mRNA decay pathway triggers degradation of most BRCA1 mRNAs bearing premature termination codons. Hum Mol Genet 11, 2805–2814.

69. Plazzer, J.P., Sijmons, R.H., Woods, M.O., Peltomäki, P., Thompson, B., Den Dunnen, J.T., and Macrae, F. (2013). The InSiGHT database: utilizing 100 years of insights into Lynch Syndrome. Familial Cancer 12, 175–180.

70. Spier, I., Yin, X., Richardson, M., Pineda, M., Laner, A., Ritter, D., Boyle, J., Mur, P., Hansen, T.V.O., Shi, X., et al. (2024). Gene-specific ACMG/AMP classification criteria for germline APC variants: Recommendations from the ClinGen InSiGHT Hereditary Colorectal Cancer/Polyposis Variant Curation Expert Panel. Genet Med 26, 100992.

71. Biesecker, L., Rehm, H., Tayoun, A.A., Berg, J., Bick, D., Byrne, A., Chao, E., Gastier-Foster, J., Karbassi, I., Moyer, A., et al. (2026). P593: Piloting the forthcoming ACMG/AMP/CAP/ClinGen standards for sequence variant classification. Genet. Med. Open 4, 104084.

72. Uzonyi, A., Dierks, D., Nir, R., Kwon, O.S., Toth, U., Barbosa, I., Burel, C., Brandis, A., Rossmanith, W., Le Hir, H., et al. (2023). Exclusion of m6A from splice-site proximal regions by the exon junction complex dictates m6A topologies and mRNA stability. Mol Cell 83, 237–251.e7.

73. Li, F., Yi, Y., Miao, Y., Long, W., Long, T., Chen, S., Cheng, W., Zou, C., Zheng, Y., Wu, X., et al. (2019). N-Methyladenosine Modulates Nonsense-Mediated mRNA Decay in Human Glioblastoma. Cancer Res 79, 5785–5798.

74. He, H., Ge, L., Chen, Y., Zhao, S., Li, Z., Zhou, X., and Li, F. (2024). m6A modification of plant virus enables host recognition by NMD factors in plants. Sci China Life Sci 67, 161–174.

75. Tan, L., Cheng, W., Liu, F., Wang, D.O., Wu, L., Cao, N., and Wang, J. (2021). Positive natural selection of N6-methyladenosine on the RNAs of processed pseudogenes. Genome Biol 22, 180.

76. Mangkalaphiban, K., Fu, L., Du, M., Thrasher, K., Keeling, K.M., Bedwell, D.M., and Jacobson, A. (2024). Extended stop codon context predicts nonsense codon readthrough efficiency in human cells. Nature Communications 15, 2486.

77. Loughran, G., Chou, M.-Y., Ivanov, I.P., Jungreis, I., Kellis, M., Kiran, A.M., Baranov, P.V., and Atkins, J.F. (2014). Evidence of efficient stop codon readthrough in four mammalian genes. Nucleic Acids Res 42, 8928–8938.

78. Ge, Z., Quek, B.L., Beemon, K.L., and Robert Hogg, J. (2016). Polypyrimidine tract binding protein 1 protects mRNAs from recognition by the nonsense-mediated mRNA decay pathway.

79. Kishor, A., Ge, Z., and Hogg, J.R. (2018). hnRNP L-dependent protection of normal mRNAs from NMD subverts quality control in B cell lymphoma. The EMBO Journal 38, EMBJ201899128.

80. Imamachi, N., Salam, K.A., Suzuki, Y., and Akimitsu, N. (2017). A GC-rich sequence feature in the 3’ UTR directs UPF1-dependent mRNA decay in mammalian cells. Genome Res 27, 407–418.

81. Hinnebusch, A.G., Ivanov, I.P., and Sonenberg, N. (2016). Translational control by 5′-untranslated regions of eukaryotic mRNAs. Science.

82. Udy, D.B., and Bradley, R.K. (2022). Nonsense-mediated mRNA decay uses complementary mechanisms to suppress mRNA and protein accumulation. Life Sci Alliance 5,.

83. Rosenfeld, N., Elowitz, M.B., and Alon, U. (2002). Negative autoregulation speeds the response times of transcription networks. J Mol Biol 323, 785–793.

84. Ni, J.Z., Grate, L., Donohue, J.P., Preston, C., Nobida, N., O’Brien, G., Shiue, L., Clark, T.A., Blume, J.E., and Ares, M., Jr (2007). Ultraconserved elements are associated with homeostatic control of splicing regulators by alternative splicing and nonsense-mediated decay. Genes Dev 21, 708–718.

85. El-Brolosy, M.A., Kontarakis, Z., Rossi, A., Kuenne, C., Günther, S., Fukuda, N., Kikhi, K., Boezio, G.L.M., Takacs, C.M., Lai, S.-L., et al. (2019). Genetic compensation triggered by mutant mRNA degradation. Nature 568, 193–197.

86. Ma, Z., Zhu, P., Shi, H., Guo, L., Zhang, Q., Chen, Y., Chen, S., Zhang, Z., Peng, J., and Chen, J. (2019). PTC-bearing mRNA elicits a genetic compensation response via Upf3a and COMPASS components. Nature 568, 259–263.

87. Mort, M., Ivanov, D., Cooper, D.N., and Chuzhanova, N.A. (2008). A meta-analysis of nonsense mutations causing human genetic disease. Hum Mutat 29, 1037–1047.

88. St Pierre, C.L., Macias-Velasco, J.F., Wayhart, J.P., Yin, L., Semenkovich, C.F., and Lawson, H.A. (2022). Genetic, epigenetic, and environmental mechanisms govern allele-specific gene expression. Genome Res 32, 1042–1057.

89. Wang, J., Chang, Y.F., Hamilton, J.I., and Wilkinson, M.F. (2002). Nonsense-associated altered splicing: a frame-dependent response distinct from nonsense-mediated decay. Mol Cell 10, 951–957.

90. Abrahams, L., Savisaar, R., Mordstein, C., Young, B., Kudla, G., and Hurst, L.D. (2021). Evidence in disease and non-disease contexts that nonsense mutations cause altered splicing via motif disruption. Nucleic Acids Res 49, 9665–9685.

91. Zetoune, A.B., Fontanière, S., Magnin, D., Anczuków, O., Buisson, M., Zhang, C.X., and Mazoyer, S. (2008). Comparison of nonsense-mediated mRNA decay efficiency in various murine tissues. BMC Genet 9, 83.

92. Nguyen, L.S., Wilkinson, M.F., and Gecz, J. (2014). Nonsense-mediated mRNA decay: inter-individual variability and human disease. Neurosci Biobehav Rev 46 Pt 2, 175–186.

93. Huang, L., and Wilkinson, M.F. (2012). Regulation of nonsense-mediated mRNA decay. Wiley Interdiscip Rev RNA 3, 807–828.

